# The Free Fatty Acid-Binding Pocket is a Conserved Hallmark in Pathogenic β-Coronavirus Spike Proteins from SARS-CoV to Omicron

**DOI:** 10.1101/2022.04.22.489083

**Authors:** Christine Toelzer, Kapil Gupta, Sathish K.N. Yadav, Lorna Hodgson, Maia Kavanagh Williamson, Dora Buzas, Ufuk Borucu, Kyle Powers, Richard Stenner, Kate Vasileiou, Frederic Garzoni, Daniel Fitzgerald, Christine Payré, Gérard Lambeau, Andrew D. Davidson, Paul Verkade, Martin Frank, Imre Berger, Christiane Schaffitzel

## Abstract

As COVID-19 persists, severe acquired respiratory syndrome coronavirus-2 (SARS-CoV-2) Variants of Concern (VOCs) emerge, accumulating spike (S) glycoprotein mutations. S receptor-binding domain (RBD) comprises a free fatty acid (FFA)-binding pocket. FFA-binding stabilizes a locked S conformation, interfering with virus infectivity. We provide evidence that the pocket is conserved in pathogenic β-coronaviruses (β-CoVs) infecting humans. SARS-CoV, MERS-CoV, SARS-CoV-2 and VOCs bind the essential FFA linoleic acid (LA), while binding is abolished by one mutation in common cold-causing HCoV-HKU1. In the SARS-CoV S structure, LA stabilizes the locked conformation while the open, infectious conformation is LA-free. Electron tomography of SARS-CoV-2 infected cells reveals that LA-treatment inhibits viral replication, resulting in fewer, deformed virions. Our results establish FFA-binding as a hallmark of pathogenic β-CoV infection and replication, highlighting potential antiviral strategies.

**One-Sentence Summary:** Free fatty acid-binding is conserved in pathogenic β-coronavirus S proteins and suppresses viral infection and replication.

## Main Text

SARS-CoV-2 causes the ongoing COVID-19 pandemic with millions of lives lost, damaging communities and economies. Human coronaviruses were previously only known to cause mild diseases of the upper respiratory tract until the emergence of the pathogenic coronaviruses SARS-CoV and Middle East respiratory syndrome coronavirus (MERS-CoV) in 2002 and 2012, respectively. Both cause severe pneumonias with a high incidence of mortality. Pathogenic SARS-CoV-2, SARS-CoV, MERS-CoV and the endemic common cold causing HCoV-OC43 and HCoV-HKU1 viruses all belong to the *beta-coronavirus* (β-CoV) genus of the *Coronaviridae* family. During the present pandemic, numerous variants of concern (VOCs) and variants of interest have emerged, exhibiting increased transmissibility, increased risk of reinfection and reduced vaccine efficiency (*1*) highlighting the urgent need for effective antiviral treatment strategies. These VOCs include the SARS-CoV-2 lineages B.1.1.7 (alpha), B.1.351 (beta), P.1 (gamma), B.1.617.2 (delta), and most recently B.1.1.529 (omicron) (*2*).

The trimeric spike glycoprotein (S) decorates the surface of coronaviruses and mediates entry into host cells. S is the major antigen recognized by neutralizing antibodies and the main target for vaccine development (*3*). SARS-CoV S and SARS-CoV-2 S both bind to human angiotensin-converting enzyme 2 (ACE2) receptor on the host cell surface (*4–6*), MERS-CoV S binds to dipeptidyl-peptidase-4 (DPP4) (*5, 7*) while the HCoV-HKU1 and HCoV-OC43 S proteins bind to the N-acetyl-9-O-acetylneuraminic acid receptor (*8*). S is cleaved by host cell proteases into the receptor-binding fragment S1 and the partially buried fusion fragment S2 (*4*). S1 is comprised of the N-terminal domain (NTD), the RBD with a receptor-binding motif (RBM), and two C-terminal domains (CTDs). S2 mediates fusion of the viral envelope with host cell membranes and is comprised of the fusion peptide, heptad repeats, transmembrane domain and cytoplasmic C-terminus (*9*). In the prefusion conformation, the RBDs in the S trimer can alternate between closed (‘down’) and open (‘up’) conformations. SARS-CoV and SARS-CoV-2 S require RBD ‘up’ conformations for interaction with ACE2 (*6, 9, 10*) for cell entry.

In our previous SARS-CoV-2 S structure, we discovered a free fatty acid (FFA) bound to a hydrophobic pocket in the RBD (*11*). Mass spectroscopy identified this ligand as LA, an essential omega-6 poly-unsaturated fatty acid (PUFA) the human body cannot synthesize (*11, 12*). LA-bound S is stabilized in a compact, locked conformation which is incompatible with ACE2 receptor binding (*11*). In immunofluorescence assays, synthetic mini-virus particles decorated with LA-bound S showed reduced docking to ACE2-expressing host cells as compared to mini-virus with LA-free S (*13*), confirming that LA interferes with receptor binding. S protein sequence alignments suggest conservation of the hydrophobic pocket in the RBDs of SARS-CoV, SARS-CoV-2, MERS-CoV, hCoV-OC43 and hCoV-HKU1 (*11*), indicating that the pocket may be a hallmark shared by all human β-CoVs. Intriguingly, all SARS-CoV-2 VOCs stringently maintain this pocket, notably including omicron which accumulated a wide range of mutations in S elsewhere, suggesting that the pocket provides a selective advantage.

Here, we investigate if LA-binding, and the functional consequences of LA-binding, is conserved in S glycoproteins of pathogenic SARS-CoV, MERS-CoV, HCoV-HKU1 as well as the SARS-CoV-2 VOCs alpha, beta, gamma, delta and omicron. We demonstrate that the respective RBDs all comprise a hydrophobic pocket capable of binding LA, except common cold-causing HCoV-HKU1. However, we also demonstrate that a single amino acid substitution of a residue lining the entrance of the hydrophobic pocket in the HCoV-HKU1 RBD is sufficient to restore LA-binding. We analyze SARS-CoV S by cryo-EM showing that LA-bound SARS-CoV S adopts a hitherto elusive locked structure akin to LA-bound locked SARS-CoV-2 S (*11*), incompatible with ACE2 receptor binding. In contrast, in the open conformation of SARS-CoV S, the pocket in the RBDs is devoid of LA. Molecular dynamics (MD) simulations corroborate spontaneous LA-binding in the respective hydrophobic pockets in the RBDs of SARS-CoV, MERS-CoV and SARS-CoV-2 VOCs, while no LA-binding to HCoV-HKU1 S is observed. Using correlative light-electron microscopy (CLEM) followed by electron tomography of SARS-CoV-2 infected cells, we provide evidence that LA, beyond counteracting infection at the S protein level, also interferes with viral replication inside infected cells. This likely occurs through inhibition of cytoplasmic phospholipase A2 (cPLA2), a key enzyme implicated in viral replication via formation of intracellular replication compartments (*14*) and in the cytokine storm causing systemic inflammation in COVID-19 (*15–17*).

LA-binding to S RBD can be analyzed by surface plasmon resonance (SPR). We previously determined a binding constant of ~41 nM for LA to SARS-CoV-2 S RBD (*11*). To corroborate our hypothesis that a functional hydrophobic pocket is evolutionarily conserved in β-CoV RBDs, we tested if RBDs from other β-CoVs also are capable of LA-binding (Fig. 1, table S1). Based on sequence alignments, the RBDs of SARS-CoV, MERS-CoV, SARS-CoV-2 and VOCs alpha, beta, gamma, delta and omicron all maintain the hydrophobic pocket, at least since the emergence of SARS-CoV in 2002 (Fig. 1A-C). In SPR experiments, LA bound with a high affinity (~72 nM) to immobilized SARS-CoV RBD (Fig. 1D), comparable to SARS-CoV-2 (*11*). Moreover, we observed a slow off-rate in agreement with tight LA-binding. LA bound with ~96 nM affinity to MERS-CoV RBD (Fig. 1E). In contrast, the RBD of HCoV-HKU1 S did not bind LA despite high sequence similarity (Fig. 1A,F). HCoV-HKU1 S comprises a bulky glutamate E375 located directly in front of the hydrophobic pocket (*18*) obstructing the pocket entrance (Fig 1F). We mutated HCoV-HKU1 E375 to alanine and restored LA-binding, albeit with a reduced affinity of ~178 nM (Fig. 1F). This indicates that the pocket function, while structurally conserved, may have been abrogated in HVoC-HKU1, a β-CoV which causes mild disease. The RBDs of SARS-CoV-2 VOCs bound LA with affinities between 50 nM and 87 nM, virtually identical to SARS-CoV-2, confirming that LA-binding is conserved (Fig. 1G) and not affected by the mutations in S which cluster away from the pocket (Fig. 1H). Taken together, we confirmed full conservation of LA-binding in highly pathogenic β-CoV S proteins, but not in S of HCoV-HKU1. Interestingly, HCoV-OC43 which likewise causes common cold, appears to also comprise a hydrophobic pocket (Fig. 1A), as seen in an earlier HCoV-OC43 S cryo-EM structure which displays unassigned density in the RBD (fig. S1) (*19*). It remains unclear what exactly this unassigned density corresponds to, which appears too small to accommodate the C18 hydrocarbon chain of LA (fig. S1).

**Fig. 1.**
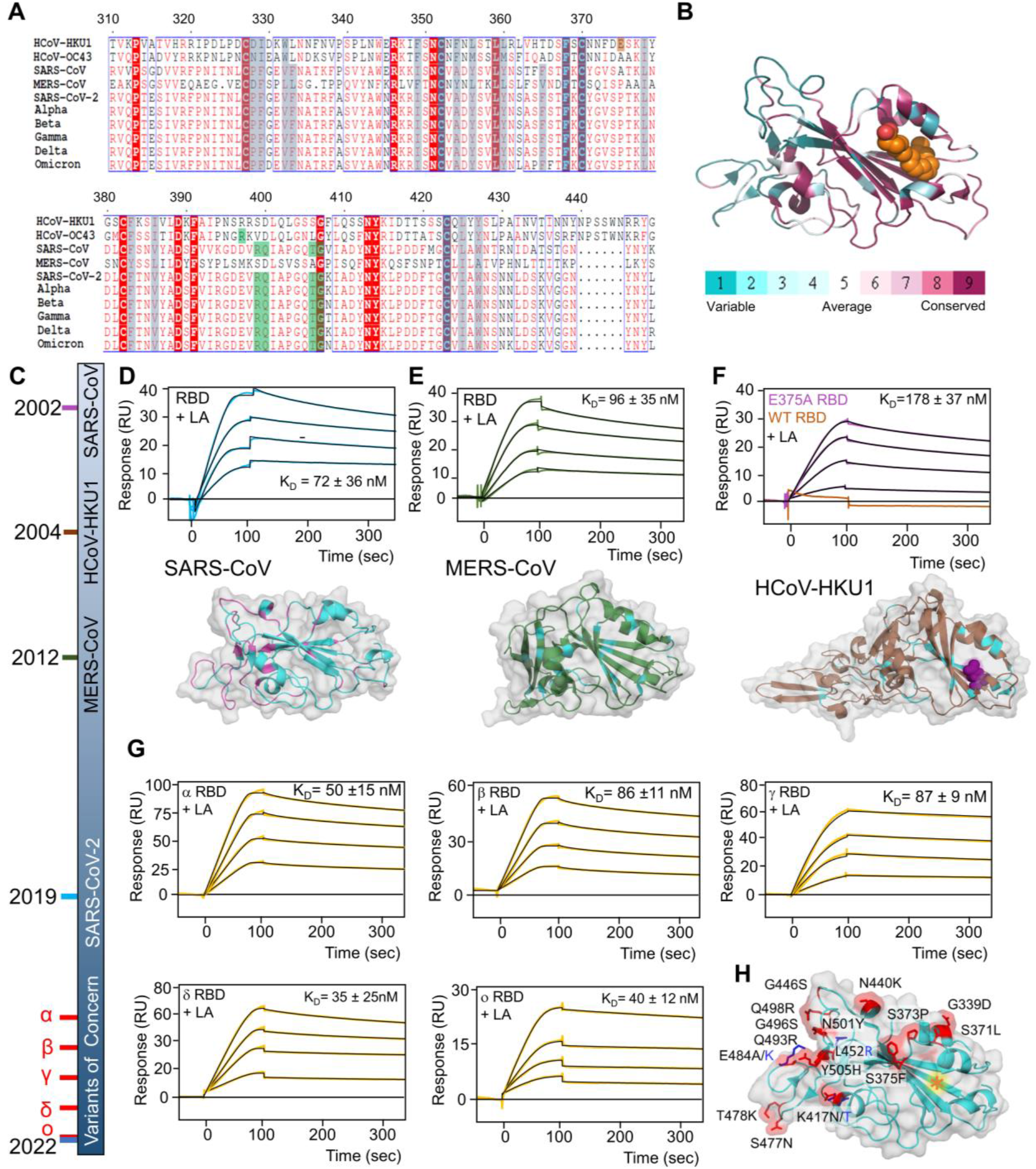
LA-binding is conserved in β-CoVs. **(A)** Alignments of the Receptor-Binding Domain (RBD) comprising the FFA-binding pocket is shown for β-CoVs HCoV-HKU1, HCoV-OC43, SARS-CoV, MERS-CoV, SARS-CoV-2 and Variants of Concern (VOCs) alpha, beta, gamma, delta and omicron. Residues lining the FFA-pocket are boxed in grey. Residues in the hydrophilic lid that interacts with the polar head group of bound FFA are highlighted in green, residue E375 in HCoV-HKU1 in orange. (**B**) Conservation of residues within the RBDs is shown, calculated with ConSurf (*34*), mapped on the structure of SARS-CoV-2 RBD (PDB ID 6ZB5). LA in the FFA-binding pocket is shown in a space-filling representation, colored in orange. (**C**) Timeline (not proportional) depicting the emergence of β-CoVs from the SARS-CoV outbreak in 2002 up to SARS-CoV-2 VOC omicron in 2022. (**D-F**) Surface Plasmon Resonance (SPR) analysis of LA-binding to immobilized RBDs of SARS-CoV (**D**), MERS-CoV (**E**), HCoV-HKU1 wild-type (WT) and E375A mutant (**F**) are shown, along with 3D structures of the respective RBDs drawn in a ribbon presentation. Residues shared with the SARS-CoV-2 RBD are colored in cyan. E375 in HCoV-HKU1 is shown in a space-filling representation, colored in purple. (**G**) LA-binding to RBDs from all five VOCs analyzed by SPR. K_D_ values are indicated. LA concentrations ranging from 4μM to 10μM were used for all SPR experiments, except for HCoV-HKU1 wildtype where only 10μM was tested. (**H**) Mutations found in the VOCs were mapped on the 3D structure of SARS-CoV-2 omicron RBD. Residue numbers and mutations are indicated. The FFA-binding pocket is marked with an orange asterisk. Mutations different from the omicron variant sequence are colored in marine blue.

To elucidate LA-binding by SARS-CoV that emerged 2002, we determined the S cryo-EM structure. The S ectodomain was produced as a secreted trimer using MultiBac (*20*) identically as described for SARS-CoV-2 S (fig. S2) (*11*). As before, we did not supplement LA during expression or subsequent sample purification and preparation steps. Cryo-EM data collection was performed with purified S protein (fig. S2, table S2). 3D classification and refinement identified an all three RBDs ‘down’ conformation and two different open conformations with one or two RBDs in the ‘up’ position, respectively (fig. S3, tables S2,S3). Using 81,242 particles, the ‘one-RBD up’ open conformation reached 3.3 Å resolution and was further analyzed (Fig. 2A, figs. S3,S4). Analysis of 178,203 particles adopting the three RBDs ‘down’ conformation yielded a 2.48 Å resolution map after applying C3 symmetry (Fig. 2A, figs. S3, S4). This three RBDs ‘down’ form of SARS-CoV S exhibits the same compact arrangement of the RBDs with fully ordered RBM, as previously identified in the LA-bound locked S structure of SARS-CoV-2 (*11*). Locked SARS-CoV S is stabilized by LA occupying a bipartite binding site composed of two adjacent RBDs in the trimer (Fig. 2B,C). One RBD contributes a ‘greasy tube’ lined by mostly phenylalanines accommodating the hydrocarbon tail of LA, and an adjacent RBD coordinates the polar headgroup of LA via residues R395 and Q396 (Fig. 2B). An overlay of the locked S structure with the previously determined LA-free closed structure of SARS-CoV S (*21*) (Fig. 2C) illustrates how LA-binding induces compaction of the three RBDs, sharing characteristic features with locked SARS-CoV-2 S (*11*). Overlay of the ‘one RBD up’ open S conformation with the closed S conformation (*21*) shows that the ‘down’ RBDs align very well (RMSD=1.086; Fig. 2D), while not aligning when overlaid with the RBDs in the locked S conformation. In agreement, we find no density for LA in the pocket of the ‘down’ RBDs of this open S conformation, and the pocket appears collapsed (Fig. 2E,F). The overlay of the RBDs from one chain in the open conformation with RBDs in the locked conformation illustrates rearrangements of Y352 and Y356 at the pocket entrance and a 5 Å movement of the gating helix to accommodate LA in the pocket (Fig. 2F). LA-binding to S also induces a profound conformational change in the loop comprising residue R620 which then engages in a cation-π interaction with Y819 (Fig. 2G). Residue Y819 is part of the fusion peptide proximal region (FPPR) of the neighboring S subunit. This conformational re-arrangement is completed by a π-π interaction of F305 and Y622 in the locked conformation (Fig. 2G). Another stabilization of the locked S trimer (and closed S trimer) is achieved by R1021 residues in each subunit which form the center of a symmetric H-bond cluster, cation-π interactions with F1024 and salt bridges to E1013 (Fig. 2H). In the open conformation this symmetry of the R1021 residue arrangement is broken, resulting in a destabilization of S (Fig. 2H). Taken together, LA-bound SARS-CoV S adopts a more stable, locked conformation incompatible with ACE2-binding, while the open conformation lacks LA and, as a consequence, is more flexible. We conclude that LA-binding is fully conserved in SARS-causing coronaviruses since at least 2002.

**Fig. 2.**
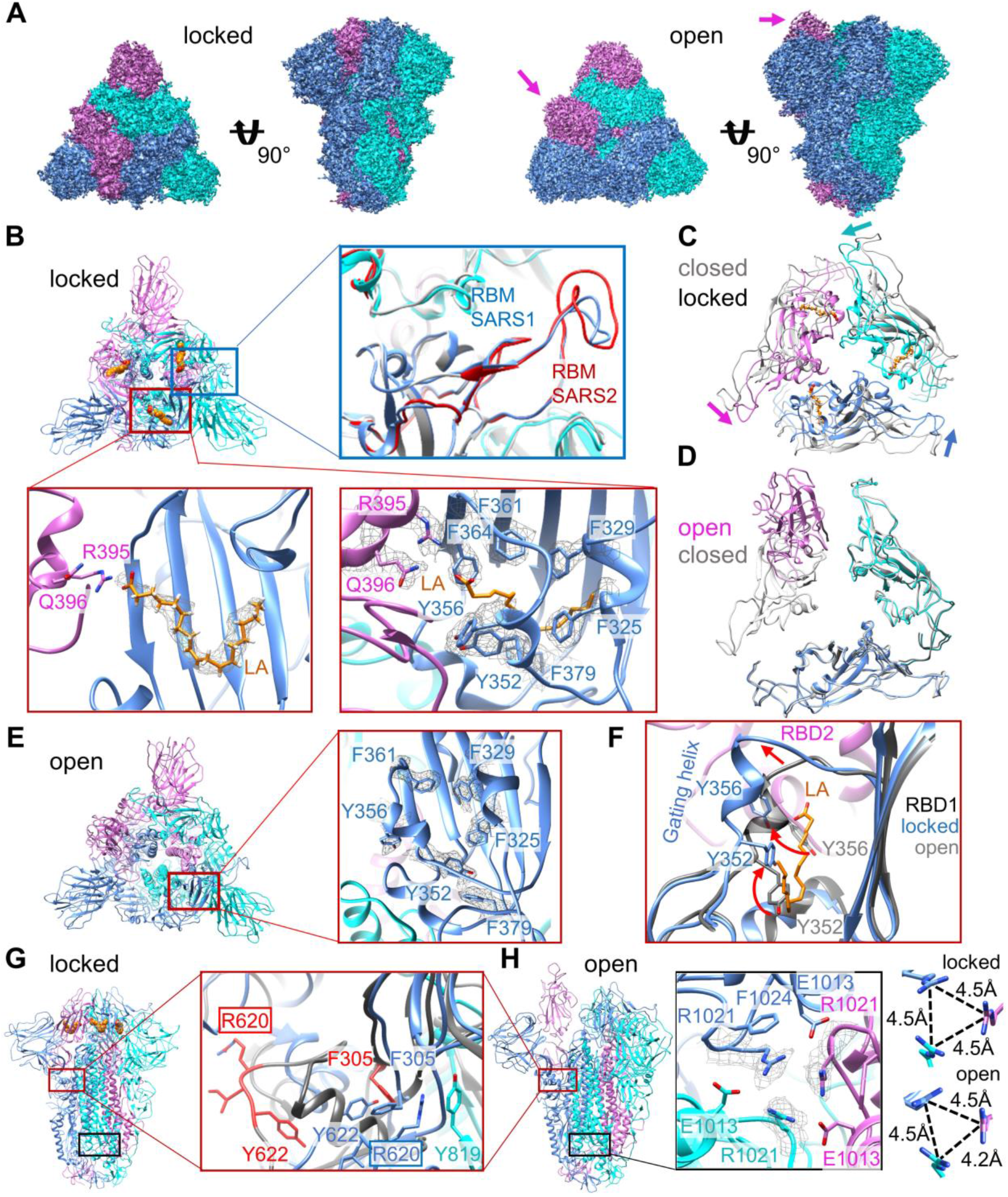
SARS-CoV S adopts an LA-bound locked conformation and an LA-free open conformation. (**A**) Cryo-EM density of SARS-CoV S trimer in the locked (left) and open (right) conformation in a front and top view. Monomers are shown in cyan, blue and magenta, respectively. Magenta arrows point to the RBD in the up conformation. (**B**) Top view of the locked S conformation in a cartoon representation. Bound LA is illustrated as spheres colored in orange. The blue box highlights the fully ordered RBM responsible for ACE2-binding, and for comparison the SARS-CoV-2 S RBM is shown in red (PDB ID 6ZB5, (*11*)). The red boxes highlight the composite LA-binding pocket formed by adjacent S RBDs. Left: the acidic LA headgroup interacts with R395 and Q396. Tube-shaped EM density is shown as mesh. Right: EM density is shown as mesh for selected hydrophobic residues lining the LA-binding pocket. (**C**) Overlay of LA-bound, locked S RBD trimers (cyan, blue, magenta) with SARS-CoV S in a closed conformation without LA ligand (grey) (PDB ID 6ACC (*21*)). (**D**) Overlay of S RBD trimers in the open conformation (cyan, blue, magenta) and the closed conformation without LA ligand (grey) (PDB ID 6ACC (*21*)). (**E**) Top view of the open conformation with one LA-binding pocket boxed in red. Right: Close-up view of the unoccupied hydrophobic pocket. EM density is shown for selected hydrophobic residues lining the entrance of the pocket (mesh). (**F**) Overlay of one subunit of LA-bound and unbound S RBD from the locked (blue) and open (grey) conformation, respectively. Red arrows show the movement of the gating helix and of Y352 and Y356 upon LA-binding. (**G**) Side view of the locked conformation. The red boxes highlight the region around R620. Close-up: The open conformation is in grey with the residues R620-Y622 and F305 in red. In the locked conformation R620 (blue) is stabilized by F305 (blue) and Y819 (cyan) from the neighboring subunit. (**H**) Side view of the open conformation. The black boxes highlight the region around R1021, forming the center of an H-bond cluster, cation-π interactions with F1024 and a salt bridge to E1013. Right: Short-range interactions formed by R1021 of S subunits have 3-fold symmetry in the locked conformation and are asymmetric in the open conformation. Distances between the carbon of the guanidinium group of the arginines (CZ atoms) are indicated.

Next, we scrutinized LA-binding to S RBDs of β-CoVs by extensive MD simulations (Fig. 3). As proof-of-principle, unbiased and spontaneous LA-binding to the SARS-CoV-2 RBD was simulated. Using the distance of the α-carbon atoms of residues N370 (gating helix) and F377 (pocket entrance) of the RBD, we monitored the dynamics of the opening and closing of the pocket during the simulations (*D_pocket*) (fig. S5A). As a measure of binding of LA in the pocket we monitored the distance between the geometric center of all carbon and oxygen atoms of LA and the center of a set of atoms that are in contact with LA when bound in the hydrophobic pocket (as determined from PDB ID 6zb5) (*D_binding*) (fig. S5B). Starting with a bound state of LA (PDB ID 6zb5 (*11*)), the system was equilibrated for 30 ns, and then LA was pushed out of the pocket by applying a small force, which is indicated by a sharp increase of the distance of LA from the center of the pocket (blue curve, Fig. 3B). Interestingly, without LA inside the pocket the distance *D_pocket* fluctuates between open and closed state (grey curve changes from 15 Å to 10 Å distance; Fig. 3B) until LA (randomly) approaches the entrance after about 600 ns, binds back to the pocket and stabilizes the open pocket (Fig.3B, movie S1). It should be noted, that during the MD simulation, LA-rebinding to the pocket was not immediate. Instead, LA transiently interacted with residues on the surface of the SARS-CoV-2 RBD (simulation time from 40 ns to 600 ns in Fig. 3B, movie S1). We identified hotspots of LA interactions on the RBD surface which include the RBM and residue N343 which is glycosylated and located close to the pocket entrance (Fig. 3A). However, spontaneous (re)binding of LA to the hydrophobic pocket was observed once LA approached the pocket entrance (Fig. 3B, fig. S5C), and LA subsequently remained stably bound in the pocket.

**Fig. 3.**
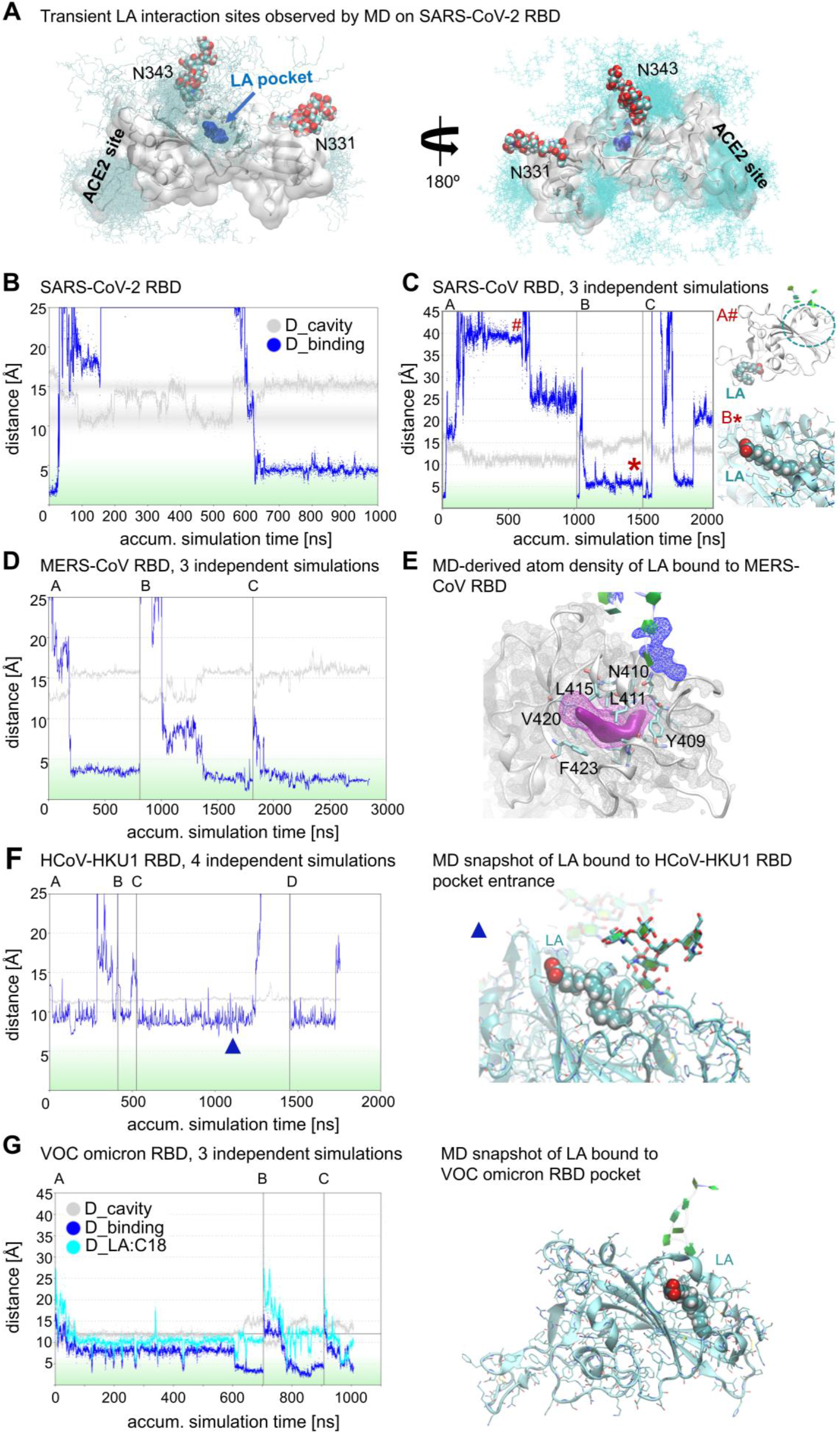
MD analysis of LA-binding to RBD of β-CoV S proteins. (**A**) LA-binding to the hydrophobic pocket of SARS-CoV-2 RBD (PDB ID 6zb5 (*11*)). LA (cyan) transiently interacts with the RBD surface before entering the pocket (dark blue). Glycosylation at residues N343 and N331 is shown as spheres. LA positions correspond to first 600 nanoseconds (ns) of MD simulation shown in panel B. Trajectories are colored identically in all panels. (**B**) Pocket dynamics (grey curve) and LA pocket binding (blue curve) to SARS-CoV-2 RBD (see also fig. S5A,B). (**C**) LA-binding to SARS-CoV RBD highlighting 3 different outcomes of simulations (A-C). Right: position of LA during the simulation (on the surface and in the pocket indicated by # and *, respectively). (**D**) LA-binding to MERS-CoV RBD, 3 simulations (A-C). (**E**) Atom-density iso-contour plot derived from accumulated 2.3 μs MD simulation of LA (magenta) bound to MERS-CoV RBD (grey ribbon and density mesh, cyan side chains, green glycosylation groups). LA high atom density is also shown as solid iso-contour volume (magenta). (**F**) LA-binding to HCoV-HKU1 RBD. Left: 4 simulation trajectories (A-D). Right: LA binds at the pocket entrance during simulation (timepoint during simulation indicated by blue triangle). (**G**) LA-binding to VOC omicron RBD. Left: 3 simulations (A-C) depicting pocket opening and binding by LA. Right: Snapshot from the end of the trajectory, with LA bound in the pocket.

Three different outcomes emerged in our MD simulations of LA (re)binding to the SARS-CoV RBD (Fig. 3C). i) LA did not (re)enter the pocket during a 1 μs simulation but interacted closely with the RBM over a significant time period (A in Fig. 3C); ii) LA rebound to the pocket after removal (B in Fig. 3C, movie S2), and iii) LA entered the pocket but could seemingly dissociate again (C in Fig. 3C). Additional simulations show that LA-binding to the RBD is dynamic because the contacts formed between LA and the residues lining the pocket vary over time and between experiments, indicating diverse binding modes (fig. S6A-D). However, when LA-binding was analyzed for the complete SARS-CoV S in the MD simulations, LA was stably bound to the three pockets formed by adjacent RBDs within the S trimer with minimal dynamics (fig. S6E,F). After validating the simulation method with experimentally derived LA-bound RBD structures (Fig. 3A-C), we applied the same MD simulation protocol to analyze LA-binding to MERS-CoV RBD (from PDB ID 6q05). Spontaneous binding of LA to the pocket of MERS-CoV RBD was observed in 10 out of 11 independent simulations (Fig. 3D,E, fig. S7A). Further analyses suggested a prevalent LA-binding mode where LA does not entirely enter the hydrophobic pocket (fig. S7B), while demonstrating significant dynamics of the portion of LA within the pocket similar to SARS-CoV (fig. S7C,D). Notably, LA-binding and pocket opening occurred simultaneously in the MERS-CoV RBD suggesting that LA can bind to the entrance of the closed pocket and pry/force the gate open (Fig. 3D, movie S3). As a control, we analyzed LA-binding to the HCoV-HKU1 RBD pocket: LA was found to transiently interact with hydrophobic residues at the pocket entrance of HCoV-HKU1 RBD but it did not enter the pocket in our simulations (Fig. 3F), reproducing our LA-binding SPR experiments (Fig. 1C). In contrast, tight LA-binding to the RBD pocket of SARS-CoV-2 VOC omicron was observed, consistent with the SPR data (Fig. 3G, movie S4).

In order to evaluate the impact of LA-treatment on SARS-CoV-2 infected cells, we infected Caco-2 cells overexpressing ACE2 (Caco-2-ACE2) with GFP-expressing SARS-CoV-2 and supplemented the cells with 50 μM LA (or solvent as a control) at 1-hour *post*-infection. Viral replication and cell viability were monitored by brightfield and fluorescence microscopy. We used CLEM followed by electron tomography to analyse SARS-CoV-2 infected Caco-2-ACE2 cells 35 hrs after LA-supplementation (Fig. 4, figs. S8-S10). Despite analysing only strong GFP-expressing cells, we detected significantly more virions in infected cells that were not LA-treated (~25 virions per micrograph) than in infected cells that had been treated with LA after infection (~9 virions per micrograph) (Fig. 4A-C, fig. S10). LA treatment also resulted in the emergence of lipid droplets in the cytoplasm of cells which appear dark in the EM micrographs (Fig. 4B, fig. S9). These droplets appear independent of whether the cells were SARS-CoV-2 infected or uninfected (Fig. 4B, figs. S9,S10), and occur in many cells including adipocytes. Moreover, we notice significant membrane remodeling in SARS-CoV-2 infected Caco-2-ACE2 cells compared to non-infected cells, as reported previously for β-CoV infections (*22, 23*) (Fig. 4, fig. S10).

**Fig. 4.**
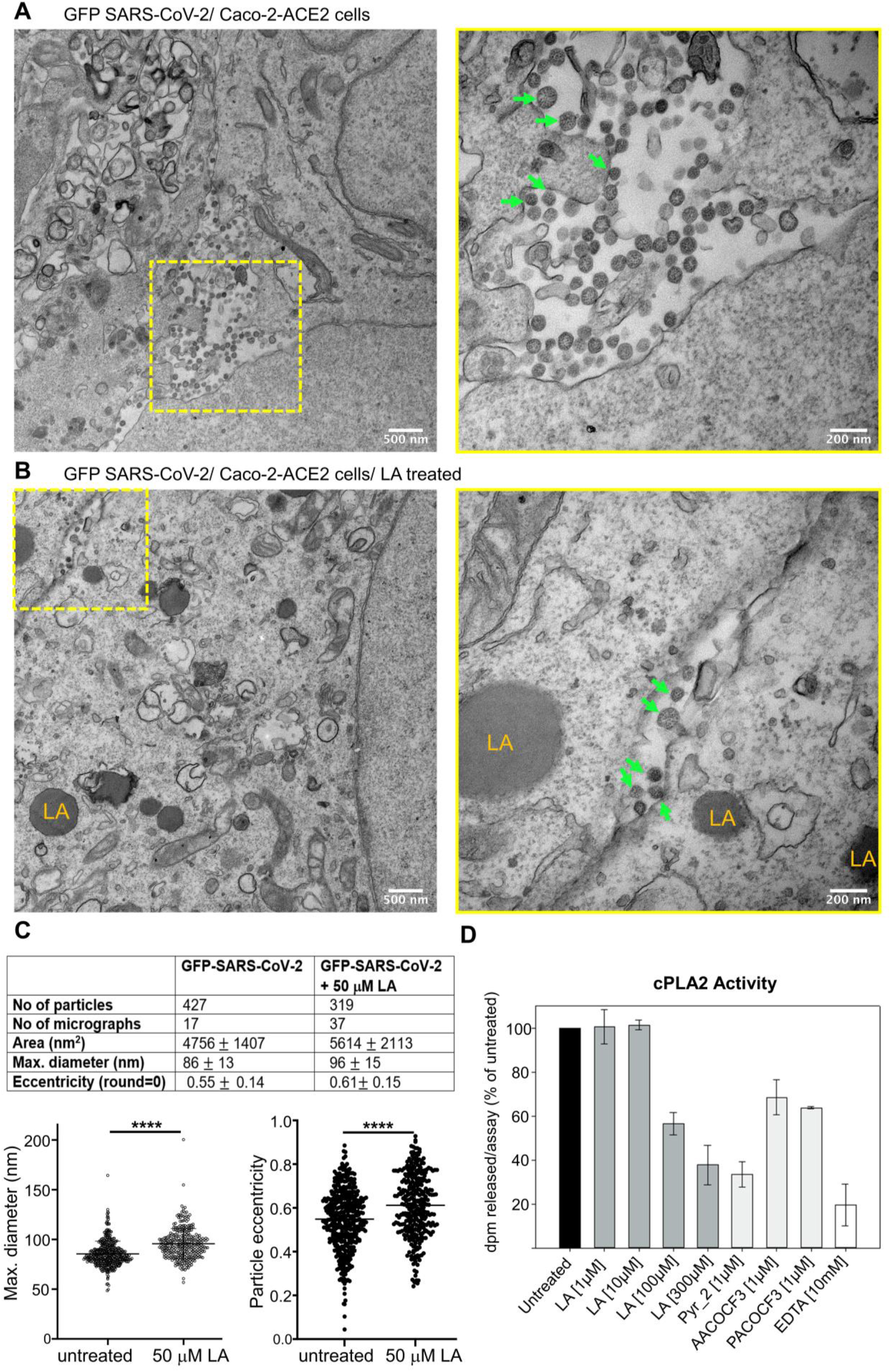
TEM analysis of GFP-expressing SARS-CoV-2 infected Caco-2-ACE2 cells. TEM images of cells in the absence **(A)** and presence **(B)** of treatment with 50 μM LA (added after infection). Left: Overview image of a GFP-SARS-CoV-2 infected cell. Yellow box highlights a region with virions. Scale bar: 500 nm. Right: Close-up view. Scale bar: 200 nm. Green arrows point to virions; LA highlights lipid droplets. (**C**) Particle quantification and size/shape analysis of virions from green cells. Values are indicated as mean ± standard deviation. Significance levels were determined by unpaired t-test. **** P < 0.0001. (**D**) Dose-response inhibition of cPLA2 by LA, compared to cPLA2 inhibitors used as positive controls (Pyr_2 (pyrrolidine-2), AACOCF3, PACOCF3 and EDTA). Data are represented as disintegration per minute (% of maximal enzymatic activity measured in the absence of inhibitor (untreated)).

In addition to their reduced number (Fig. 4C), virions in SARS-CoV-2 infected, LA-treated cells appeared irregular in size and shape as compared to the virions in untreated cells which adopt a characteristic regular spherical form (Fig. 4A,B, fig. S10, movies S5, S6). Closer analysis of virions derived from LA-treated cells confirmed a statistically significant increase in size and deformation (ellipticity) (Fig. 4C). The average diameter of virions from untreated cells (n=427) was calculated as 86 nm (SD=13 nm), in agreement with previous reports (*14*). In contrast, virions derived from LA-treated cells (n=319) had a larger average diameter of 96 nm (SD=15 nm). Consistently, the average area of virions increased from 4756 nm^2^ to 5614 nm^2^ (Fig. 4C). Moreover, the average ellipticity of virions increased from 0.55 (SD=0.14) in untreated cells to 0.61 (SD=0.15) in LA-treated cells, confirming deformation. Taken together, we observed that LA treatment after infection leads to a lower viral load in SARS-CoV-2 infected cells with the virions being larger in size and deformed as compared to untreated, infected cells (Fig. 4A-C) indicating that their integrity, and potentially their infectivity, may be compromised.

During early stages of coronavirus infection, phospholipase A2s (PLA2s) are activated as evidenced by increased cellular levels of lysophospholipids and FFAs such as LA, arachidonic acid, oleic acid and palmitoleic acid in both SARS-CoV-2 infected cell cultures and SARS-CoV-2 infected patients (*16, 24–26*). A central regulator of lipidome remodeling during β-CoV infection is cytoplasmic PLA2 (cPLA2). Inhibition of this enzyme interferes with coronavirus-induced membrane rearrangements including formation of intracellular double-membrane vesicles and replicative organelles, which are essential for viral replication (*24*). cPLA2 cleaves glycerophospholipids at the sn-2 ester position, generating FFAs and lysophospholipids. Previously, it was shown that cPLA2 is tightly regulated by PUFAs, including LA, which are potent competitive inhibitors of the enzyme (*27*). We find that cPLA2 inhibition is half-maximal in the presence of ~100 μM LA *in vitro* (Fig. 4D). We note that the S proteins of pathogenic β-CoVs have orders of magnitude higher affinity for LA (Fig. 1). It is thus likely that, by utilizing FAs and glycerophospholipids for building viral membranes in cells, and by direct LA-binding to S, pathogenic β-CoVs overcome cPLA2 feedback inhibition to enhance membrane remodeling during infection.

Taken together, the remarkably conserved FFA-binding pocket emerges as a hallmark of pathogenic β-CoV S structure, consistent with essential functions beyond ligand-binding. The cryo-EM structure of SARS-CoV S demonstrates stringent conservation of the LA-binding pocket since at least 2002 when the SARS-CoV outbreak occurred (Fig. 2). We observe a high correlation between LA-binding in the pocket and the locked conformation which is a non-infectious form of S, incompatible with ACE2 receptor binding (Fig. 2B). MD simulations consistently show transient interactions of LA with hydrophobic patches on the surface of the RBDs before binding the pocket, suggesting a conserved mechanism of LA approaching the pocket entrance (Fig. 3). However, stable LA-binding to the RBDs only occurs once LA has entered the pocket. It is less clear if the pocket first has to open by moving the gating helix, or if LA-binding at the entrance of the pocket can pry open the pocket. We find examples for both scenarios in the different simulations, but an opening of the pocket while interacting with LA (induced fit) occurred more frequently (Fig. 3B,D).

Previous studies showed that LA renders the virus less infectious by stabilizing a locked form of S inhibiting receptor-binding (*11*), virion attachment and entry into cells (*13*). Here we show that LA treatment of infected cells significantly reduces the production of virions, and the few virions produced are markedly deformed (Fig. 4A-C, fig. S10). These modes of action will likely synergize to significantly reduce, or even abrogate, viral infectivity and transmission. Therefore, FFA-binding by S can be conceived as an ‘Achilles heel’ of pathogenic β-CoVs, making this feature a highly attractive target for LA- or LA-mimetic based antiviral interventions against SARS-CoV-2. It is noteworthy that fatty acids have been used prolifically as excipients in unrelated medications with established safety in multiple administration routes (nasal, pulmonary, oral and intra-venous) (*28*).

The evolutionary conservation of the FFA-pocket implies significant selection advantage for the virus itself. We can conceive of several such advantages. For instance, the LA-bound S form is more stable than the open S forms (*29*). LA stabilizes the S protein in a prefusion conformation which is particularly useful when the protein is still in the host cell, preventing premature proteolytic cleavage by host proteases. Moreover, LA-bound, locked S buries key epitopes of the RBM and of core RBD parts, potentially hiding these from attack by neutralizing antibodies (*1, 13*). We speculate that LA levels can be sensed by the virus, allowing the S protein to switch from the ‘stealth’ locked form to the infectious open form that mediates ACE2-binding and cell entry. Lipid metabolome remodeling is a central element of viral infection (*25, 30, 31*). LA levels are markedly perturbed during COVID-19 disease progression, with serum levels significantly decreased in COVID-19 patients (*32*). Conversely, intracellular levels of LA are elevated (*25*). This correlates with β-CoV-induced membrane remodeling to generate new membrane compartments for viral replication (*22, 23*). cPLA2 activation is a central mediator of lipidome remodeling in β-CoVs, and of various additional +RNA virus families, and is therefore a validated target for broad-spectrum antiviral drug development (*24*). We propose that during β-CoV infection when cPLA2 is activated, the virus can circumvent feedback inhibition of cPLA2 by sequestering LA (*27*) (Fig. 4D). This would keep cPLA2 in a hyperactivated state. In this model, inhibition of cPLA2 by supplementing excess LA will downregulate membrane remodeling, thus interfering with a mechanism required for viral replication. Indeed, we demonstrate that supplementing excess LA to SARS-CoV-2 infected cells interferes with viral replication (Fig. 4). Previous reports show that LA also interferes with MERS-CoV replication (*25*). In conclusion, our results convey that the conserved FFA-S interaction, while affording selective advantages to the virus, renders it vulnerable to antiviral intervention exploiting this highly conserved feature. This could be achieved by supplementing LA or a related molecule, ideally during early stages of infection when the virus resides in the upper respiratory tract where it can be conveniently targeted. This could be accomplished by delivering FFA formulations, *e.g*. via a nasal or inhaled spray, to suppress viral replication and spreading within a patient, concomitantly reducing transmission (*33*) and protecting others from infection.

## Acknowledgments

We thank all members of the Berger and Schaffitzel laboratories for their contributions and suggestions, and David Veesler (University of Washington, Seattle) and Roman Laskowski (EMBL-EBI) for helpful discussions. We acknowledge support and assistance by the Wolfson Bioimaging Facility and the GW4 Facility for High-Resolution Electron Cryo-Microscopy funded by the Wellcome Trust (202904/Z/16/Z and 206181/Z/17/Z) and BBSRC (BB/R000484/1). We are grateful for support from the Oracle Higher Education and Research program to enable cryo-EM data processing using Oracle’s high-performance public cloud infrastructure (https://cloud.oracle.com/en_US/cloud-infrastructure), and we thank Simon Burbidge, Thomas Batstone and Matt Williams for computation infrastructure support. We are grateful for computing time sponsored by BIOGNOS AB, Göteborg. The authors are grateful to University of Bristol’s Alumni and Friends, which funded the ImageXpress Pico Imaging System. I.B. acknowledges support from the European Research Council (ERC AdvG DNA-DOCK) and the EPSRC Future Vaccine Manufacturing and Research Hub (EP/R013764/1). C.S. and I.B. are Investigators of the Wellcome Trust (210701/Z/18/Z; 106115/Z/14/Z). M.K.W. was supported by MRC grant MR/V027506/1 (awarded to A.D.D). G.L. acknowledges support from the Fondation Jean Valade/Fondation de France (Award FJV_FDF-00112090), the National Research Agency (AirMN (ANR-20-CE14-0024-01)) and “Investments for the Future” Laboratory of Excellence SIGNALIFE, a network for innovation on signal transduction pathways in life sciences (ANR-11-LABX-0028-01 and ANR-15-IDEX-01), and the Fondation de la Recherche Médicale (DEQ20180339193L). Recombinant human cPLA2 was a kind gift from Pr. Michael H. Gelb, University of Washington, Seattle, USA.

## Author contributions

C.S. and I.B. conceived and guided the study. K.G., F.G. and K.V. produced sample. K.G. carried out SPR experiments. S.K.N.Y. and U.B. prepared grids, collected and processed EM data. C.T., D.B. and K.P. carried out image analysis, model building and refinement. C.T. interpreted structures. M.F. performed and analyzed all MD simulations. A.D.D. and M.K.W. performed all live virus CL3 work and light microscopy. L.H. and P.V. performed EM and interpreted the image/tomograms. C.P. and G.L., assisted by R.S., performed the cPLA2/LA enzymatic assays. C.T., K.G., D.F., I.B. and C.S interpreted results. C.S. and I.B. wrote the manuscript with input from all authors.

## Competing interests

The authors declare competing interests. I.B. and F.G. are shareholders of Imophoron Ltd, unrelated to this correspondence. D.F. and I.B. are shareholders of Geneva Biotech SARL, unrelated to this correspondence. I.B., C.S. and D.F. report shareholding in Halo Therapeutics Ltd related to this Correspondence. Patents and patent applications have been filed related to FFA therapeutic interventions. The other authors do not declare competing interests.

## Data and materials availability

All datasets generated during the current study have been deposited in the Electron Microscopy Data Bank (EMDB) under accession numbers EMD-14718 (C1 locked conformation), EMD-14717 (C3 locked conformation) and EMD-14724 (RBD-up open conformation), and in the Protein Data Bank (PDB) under accession numbers 7ZH2 (C1 locked conformation), 7ZH1 (C3 locked conformation) and 7ZH5 (RBD-up open conformation). Reagents are available from I.B. and C.S. under material transfer agreement with the University of Bristol.

## Supplementary Materials

Materials and Methods

Figs. S1 to S10

Tables S1 to S3

References (*35*–*59*)

Movies S1 to S6

