## Supplementary Data for "The Free Fatty Acid-Binding Pocket is a Conserved Hallmark in Pathogenic β-Coronavirus Spike Proteins from SARS-CoV to Omicron"

**This PDF file includes:**

Materials and Methods

Figs. S1 to S10

Tables S1 to S3

References (*35–59*)

Captions to Movies S1 to S6

**Other Supplementary Material for this manuscript includes the following:**

Movies S1 to S6

### Materials and Methods

Protein production

*SARS-CoV S protein.* The cDNA for expression of SARS-CoV S ectodomain (Uniprot ID-P59594, Isolate BJ01) was codon-optimized for insect cell expression and synthesized by Genscript (Genscript Inc, New Jersey USA). In this construct, S ectodomain comprises amino acids 14 to 1193, preceded by the GP64 secretion signal sequence (amino acid sequence MVSAIVLYVLLAAAAHSAFA) at the N-terminus. The construct is fused to a C-terminal thrombin cleavage site followed by a T4-foldon trimerization domain and a hexa-histidine affinity purification tag. The protein was expressed using the MultiBac baculovirus expression system (Geneva Biotech, Geneva, Switzerland) (*20*) in Hi5 cells using ESF921 media (Expression Systems Inc.). Three days post-transfection, the supernatant from transfected cells was harvested by centrifugation at 1,000g for 10 min followed by a second centrifugation of the supernatant at 5,000g for 30 min. The final supernatant was incubated with 5 ml HisPur Ni-NTA Superflow Agarose (Thermo Fisher Scientific) per 2 liters of culture overnight at 4°C. A gravity flow column was used to collect the resin bound with SARS-CoV S protein. The resin was washed extensively with wash buffer (65 mM $NaH_2PO_4$, 300 mM NaCl, 20 mM imidazole, pH 7.5), and the protein was eluted using elution buffer (65 mM $NaH_2PO_4$, 300 mM NaCl, 235 mM imidazole, pH 7.5). Elution fractions were analyzed by reducing SDS-PAGE and fractions containing SARS-CoV S protein were pooled and concentrated using 50 kDa MWCO Amicon centrifugal filter units (EMD Millipore) and buffer-exchanged in SEC buffer (20 mM Tris, pH 7.5, 100 mM NaCl). Concentrated SARS-CoV S was subjected to size exclusion chromatography (SEC) using a Superdex 200 increase 10/300 column (GE Healthcare) in SEC buffer. Peak fractions from SEC were analyzed by reducing SDS-PAGE and negative stain electron microscopy (EM). Fraction 8 was used for cryo-EM (fig. S2).

*Biotinylated SARS-CoV Receptor-Binding Domain (RBD).* The SARS-CoV RBD-encoding DNA was codon-optimized for insect cell expression and synthesized by Genscript (Genscript Inc., New Jersey USA). This construct comprised amino acid residues 306 to 527 and was fused at its N-terminus to the SARS-CoV-2 S secretion signal sequence (amino acid sequence MFVFLVLLPLVSSQ) and was followed by a linker (amino acid sequence GGSGGSGSG), an avi-tag (amino acid sequence GLNDIFEAQKIEWHE), a second linker (amino acid sequence GSGSGS) and finally an octa-histidine tag for purification. This construct was inserted into pACEBac1 plasmid (Geneva Biotech, Geneva, Switzerland). The protein was produced and purified as described above for SARS-CoV S ectodomain except the last SEC step, which was performed in 1x phosphate buffered saline pH 7.5 (PBS). Biotinylation was achieved by incubation with BirA in the presence of biotin according to established protocols (*35*). Remaining free biotin and BirA were removed by purifying biotinylated SARS-CoV RBD by SEC using a S200 10/300 increase column (GE Healthcare).

*Biotinylated MERS-CoV Receptor-Binding Domain (RBD).* The biotinylated MERS-CoV RBD expression construct was generated as described above for biotinylated SARS-CoV RBD. The construct comprises MERS-CoV (Uniprot ID- K0BRG7) residues 367 to 606. The protein was expressed and purified as described above.

*Biotinylated HCoV-HKU1 Receptor-Binding Domain (RBD) (wild-type and E375A mutant).* Biotinylated HCoV-HKU1 RBD expression constructs were generated as described above for biotinylated SARS-CoV RBD. The wild-type construct contains HCoV-HKU1 (Uniprot ID-U3N885) residues 310 to 624. In E375A glutamate 375 was mutated to alanine. The wild-type and E375A RBD proteins were expressed and purified as described above.

*Biotinylated SARS-CoV-2 VOC Receptor-Binding Domains (RBD).* Biotinylated *SARS-CoV*-2 RBDs from variant expression constructs were generated as described above for biotinylated SARS-CoV RBD. These constructs contain SARS-CoV-2 (Uniprot ID- P0DTC2) residues 319 to 541, with variant specific mutations. The RBDs were expressed and purified as described above.

### Negative-stain sample preparation and electron microscopy

4 μL of 0.05 mg/mL SARS-CoV S protein was applied onto a freshly glow discharged (1 min at 10 mA) CF300-Cu-50 grid (Electron Microscopy Sciences), incubated for 1 min, and manually blotted. 4 μL of 3% Uranyl Acetate was applied onto the same grid and incubated for 1 min before the solution was blotted off. Images were acquired at a nominal magnification of 49,000x on a FEI Tecnai 12 120 kV BioTwin Spirit microscope.

### Cryo-EM sample preparation and data collection

4 μL of 0.05 mg/mL SARS-CoV S protein was loaded onto a freshly glow discharged (2 min at 4 mA) Quantifoil R1.2/1.3 carbon grid (Agar Scientific), blotted using a Vitrobot MarkIV (Thermo Fisher Scientific) at 100% humidity and 4°C for 2s, and plunge frozen. Data were acquired on a FEI Talos Arctica transmission electron microscope operated at 200 kV and equipped with a Gatan K2 Summit direct detector and Gatan Quantum GIF energy filter, operated in zero-loss mode with a slit width of 20 eV using the EPU software.

Data were collected in counted super-resolution mode at a nominal magnification of 130,000x with a physical pixel size of 1.05 Å/pix and a virtual pixel size of 0.525 Å/pix. The dose rate was adjusted to 5.77 counts/physical pixel/s. Each movie was fractionated in 60 frames of 200 ms. 6,600 micrographs were collected with a defocus range comprised between -0.8 and -2.0 µm.

### Cryo-EM data processing

The dose-fractionated movies were gain-normalized, aligned, and dose-weighted using MotionCor2 (*36*). Defocus values were estimated and corrected using the Gctf program (*37*). 1,724,689 particles were automatically picked using Relion 3.0 (*38*). Reference-free 2D classification was performed to select well-defined particles. After three rounds of 2D classification, a total of 784,580 particles were selected for further 3D classification. The initial 3D model (*11*) was filtered to 60 Å during 3D classification in Relion using 8 classes. Classes 4 and 5 (fig. S3), showing prominent features of closed conformation representing a total of 178,203 particles were combined and used for 3D refinement. Class 6 comprised S proteins in a one RBD up (open) conformation, and Class 1 presented S in a two RBD up (open) conformation, comprising 81,707 and 122,315 particles respectively (fig. S3). The selected maps were subjected to 3D refinement without applying any symmetry. Subsequently, the maps were subjected to local defocus correction and Bayesian particle polishing in Relion 3.1. Global resolution and B factor (-68 $Å^2$ and -100.8 $Å^2$ for closed and open maps respectively) of the maps were estimated by applying a soft mask around the protein density, using the gold-standard Fourier Shell Correlation (FSC) criterion 0.143, resulting in an overall resolution of 2.71 Å and 3.34 Å respectively (fig. S4). C3 symmetry was applied to the closed conformation map using Relion 3.1, followed by CTF refinement and Bayesian polishing, yielding a final resolution of 2.48 Å (B factor of -73.71 $Å^2$) (fig. S4). Local resolution maps were generated using Relion 3.1 (fig. S4).

Cryo-EM model building and analysis

UCSF Chimera (*39*) was used to fit atomic models of the SARS-CoV S closed conformation (PDB ID 6ACC (*21*)) and open conformation (PDB ID 6ACD (*21*)) into our SARS-CoV C1 closed conformation and open conformation cryo-EM map respectively. To improve the model building we used the symmetrized C3 map for the closed conformation, as well as Namdinator (*40*) for open and closed conformations. Model building was done in Coot (*41*) with unsharpened and sharpened maps (*42*) and N-linked glycans were built into the density for all three models where visible (table S3). The RBD-up in the open conformation was fitted as a rigid body into the corresponding density because the resolution in this part of the EM structure is not sufficient to build an atomic model. Restraints for the Linoleic Acid (LA) were generated with eLBOW (*43*). The models for C1 and C3-symmetrized closed conformation and the open conformation were real space refined with Phenix (*44*), and the quality was checked using MolProbity (*45*) and EMRinger (*46*). Figures were prepared using UCSF chimera and PyMOL (Schrodinger, Inc).

Surface plasmon resonance (SPR) experiments

Interaction experiments using surface plasmon resonance (SPR) between LA and different RBDs were carried out with a Biacore T200 system (GE Healthcare) according to the manufacturer's protocols and recommendations and as described previously (*11*). Briefly, biotinylated RBDs were immobilized on streptavidin-coated SA sensor chips at ~2500 RUs. LA sodium salt was dissolved in PBS pH 7.5 at a concentration of 10 mM and then serially diluted and injected at concentrations of 4 mM, 6 mM, 8 mM and 10 mM at a flowrate of 30 µl/minute. The running buffer for all SPR measurements was PBS buffer pH 7.5. The sensorgrams were analyzed using the Biacore Evaluation Software (GE Healthcare) and $k_{on}$, $k_{off}$ and $K_D$ values were determined by fitting the raw data individually for each concentration using a 1:1 binding model. All experiments were performed in triplicates.

Molecular Dynamics Simulations

Starting structures for the simulations were setup using the graphical interface of YASARA (*47*). For SARS-CoV-2 RBD (residues 319-592) PDB ID 6zb5 (pocket open state, LA-bound) (*11*) was used. Starting structures of SARS-CoV RBD (residues 307-577) were built based on the structure reported here (pocket open state, LA-bound). MERS-CoV RBD (residues 368-655) was modelled based on PDB IDs 6q05 or 4l3n (pocket closed state) (*48, 49*). PDB ID 5gnb was used for HCoV-KHU1 RBD (residues 311-674) (*18*).

In general, the systems were solvated in 0.9% NaCl solution (150 mM), and simulations were performed at 310 K using periodic boundary conditions and using the AMBER14 force field. The box size was rescaled dynamically to maintain a water density of 0.996 g/ml. Simulations were performed using YASARA with GPU acceleration in 'fast mode' (4 fs time step) (*50*) on 'standard computing boxes', e.g. equipped with one 12-core i9 CPU and NVIDIA GeForce GTX 1080 Ti.

Three types of molecular simulation protocols were performed, termed here "LA bound", "LA unbinding" and "LA *de novo* binding". In the first and second protocol the starting structure consists of an RBD/LA complex that has the LA molecule bound in the experimentally determined binding pocket ('open pocket'). "LA bound" follows a standard NPT MD protocol. The second MD protocol ("LA unbinding") consists of three periods: about 30 ns equilibration of the bound state, a short period where the LA was pushed out of the binding pocket by application of a force of 1 kcal/mol, finally followed by a long MD sampling period in which the LA was free to diffuse through the bulk solvent or interact with the protein surface. The third MD protocol ("LA *de novo* binding") aims to simulate LA binding '*de novo*' based on an experimentally determined RBD 'apo' structure with a hidden LA pocket ('closed pocket') and LA molecule(s) positioned in the bulk solvent. Although binding events of drug fragments have been simulated previously on special-purpose supercomputers designed specifically for MD simulations (*51*), the unbiased simulation of small molecule binding events can currently be still considered a 'challenging endeavor' in computational chemistry.

During this project, in total more than 150 MD trajectories were sampled starting with LA either in bound or unbound to RBD state: 36 for SARS-CoV-2 WT (15 µs accumulated timescale), 9 for SARS-CoV-2 K417N, E484K, N501Y (5 µs accumulated timescale), 74 for SARS-CoV (20 µs accumulated timescale), 32 for MERS-CoV (11 µs accumulated timescale), 9 for HCoV-KHU1(4 µs accumulated timescale). Only scientific plots of the most relevant trajectories are shown in order to keep the complexity of the data presented reasonable. Further details can be found in the captions of the figures S5-S7.

Conformational Analysis Tools (CAT, http://www.md-simulations.de/CAT/) was used for analysis of trajectory data, general data processing and generation of scientific plots. VMD (*52*) was used to generate molecular graphics.

### Live SARS-CoV-2 experiments

*Cells and virus propagation.* A VeroE6 cell line modified to constitutively express the serine protease TMPRSS2 (Vero E6/TMPRSS2, obtained from NIBSC, UK) and the human gut epithelial cell line Caco2 expressing ACE2 (Caco-2-ACE2, a kind gift of Dr Yohei Yamauchi, University of Bristol) were cultured at 37 °C in 5% $CO_2$ in Dulbecco's modified Eagle's medium plus GlutaMAX (DMEM, Gibco, ThermoFisher) supplemented with 10% fetal bovine serum (FBS, Gibco, ThermoFisher) and 0.1 mM non-essential amino acids (NEAA, Sigma Aldrich). A SARS-CoV-2 reporter virus expressing a gene encoding the fluorescent protein turboGFP in place of the ORF7 gene (termed rSARS-CoV-2/Wuhan/ORF7-tGFP) was generated using a SARS-CoV-2 (Wuhan isolate) reverse genetics system utilizing the "transformation-associated recombination in yeast" approach (*53*). 11 cDNA fragments with 70 bp end-terminal overlaps which spanned the entire SARS-CoV-2 isolate Wuhan-Hu-1 genome (GenBank accession: NC_045512) and replaced ORF7 gene with the turboGFP gene were produced by GeneArt™ synthesis (Invitrogen™, ThermoFisher) as inserts in sequence verified, stable plasmid clones. The 5´-terminal cDNA fragment was modified to contain a T7 RNA polymerase promoter and an extra "G" nucleotide immediately upstream of the SARS-CoV-2 5´ sequence, whilst the 3´-terminal cDNA fragment was modified such that the 3' end of the SARS-CoV-2 genome was followed by a stretch of 33 "A"s followed by the unique restriction enzyme site *Asc*I. The inserts were amplified by PCR using a Platinum SuperFi II mastermix (ThermoFisher) and assembled into a full-length SARS-CoV-2 cDNA clone in the YAC vector pYESL1 using a GeneArt™ High-Order Genetic Assembly System (A13285, Invitrogen™, ThermoFisher) according to the manufacturer's instructions. RNA transcripts produced from the YAC clone by transcription with T7 polymerase were used to recover infectious virus. Whole genome sequencing confirmed the virus sequence. The virus was propagated in VeroE6/TMPRSS cells grown in infection medium (Eagle's minimum essential medium plus GlutaMAX (MEM, Gibco) supplemented with 2% FBS and NEAA). Cells were incubated at 37 ºC in 5% $CO_2$ until cytopathic effects were observed at which time the supernatant was harvested and filtered through a 0.2µm filter, aliquoted and stored at -80 °C.

*Viral detection by fluorescence.* Caco-2-ACE2 cells were seeded onto 9 mm glass coverslips coated in a finder pattern of evaporated carbon (~10 nm) and poly-D-lysine in 24 well plates or in µClear 96-well Microplates (Greiner Bio-one) in DMEM supplemented with 10% FBS until cell coverage on the coverslips reached 25%. The cells were inoculated with rSARS-CoV-2/Wuhan/ORF7-tGFP at a multiplicity of infection (MOI) of 5 in infection medium for 60 minutes at room temperature before the media was removed and replaced with infection medium containing 50 µM LA and 0.25% DMSO, or 0.25% DMSO only. Control wells were treated the same but received no infectious inoculum. Cells were incubated at 37 °C in 5% $CO_2$ for 36 hours until turboGFP expression was detectable in cells in the 96 well plate by fluorescence imaging with an ImageXpress Pico Automated Cell Imaging System (Molecular Devices). Samples were inactivated and fixed by submersion in 4% paraformaldehyde (PFA) for 60 minutes at room temperature. All work with infectious recombinant SARS-CoV-2 was done inside a class III microbiological safety cabinet in a containment level 3 facility at the University of Bristol.

*Immunofluorescence analysis.* Fixed coverslips were stained with 1mg/ml of DAPI for 5 min and transferred to a 24-well imaging plate with an ultrathin (25 mm) film bottom (Eppendorf) containing 500 µl of PBS. Images were acquired on a Leica SP5II AOBs confocal laser scanning microscope attached to an inverted DMI600 epifluorescence microscope using a 10x dry objective (0.3NA) and a 63x oil-immersion objective (1.4NA). Low magnification overviews of coverslips were acquired to identify location of region of interest on carbon finder pattern.

*TEM Sample preparation.* Following fluorescence imaging, coverslips were rinsed in 0.1 M sodium cacodylate buffer (pH 7.4) and post-fixed in 2.5% glutaraldehyde in 0.1 M sodium cacodylate buffer at 4°C until further processing. Samples were subsequently stained and further cross-linked with osmium-ferrocyanide (1% OsO4, 1.5 % $K_4Fe(CN)_6.3H_2O$, 0.1M sodium cacodylate buffer) for 1 hour at 4°C, before en bloc staining in 3% uranyl acetate for 30 min. Following a dehydration series at room temperature in ethanol (70%, 80%, 90%, 96%, 100%), coverslips were infiltrated with 50% epoxy resin (Agar Scientific) in propylene oxide for 1 hour, followed by 100% epoxy resin 2 times for 30 min. Coverslips with cells facing up were covered in fresh epoxy resin and polymerized at 60°C for 48 hours. After ~16 hours an epoxy resin stub was placed on top of the coverslip and the samples returned to the oven. Coverslips were removed from blocks using liquid nitrogen and boiling $H_2O$ to reveal the carbon finder pattern. The blocks were trimmed to the region of interest and sectioned using a UC6 Leica ultramicrotome with a diamond knife (Diatome). Ultrathin sections (70 nm) were collected onto pioloform-coated slot grids and post stained with uranyl acetate and lead citrate for 10 min and 4 min respectively before imaging at 120KV using a Tecnai 12 BioTwin Spirit TEM. Virus particles were measured in images acquired at 18,500 x magnification in FIJI, as described (*54*).

*Electron tomography*. 300 nm sections collected on Pioloform-coated slot grids (Agar Scientific) were incubated in a solution of 15 nm gold fiducial markers (Aurion) for 5 min on each side. Tilt series (-65° to +65° at 1.5° increments) were acquired at 19,000 x magnification (0.5261 nm/px) using a FEI Tecnai 20 transmission electron microscope operated at 200 kV and equipped with a 4k-by-4k FEI Eagle camera. Electron tomograms were reconstructed using fiducial markers for alignment in IMOD (*55*).

cPLA2 Activity Assays

Recombinant human cytosolic PLA2 (cPLA2 group IVA, PLA2G4A) was expressed and purified from insect cells as described (*56*). The enzymatic activity of cPLA2 was measured using *E. coli* membranes radiolabeled with [$^3$H]-oleic acid as described (*57*). To measure the inhibitory effect of LA, recombinant human cPLA2 (10 nM final concentration) was preincubated in the absence or presence of various concentrations of LA (1 to 300 µM) for 15 min in 100 µl of PLA2 activity buffer (100 mM Tris-HCl pH 8.0, 10 mM $CaCl_2$, and 0.1% BSA) after which 30,000 dpm of radiolabeled *E. coli* membranes was added (diluted in 100 µl of buffer), with further incubation for 1 hour at 37 °C. Enzymatic reactions were stopped by addition of 1 volume (200 µl) of PLA2 stop buffer (100 mM EDTA, 0.2% fatty acid-free BSA). Tubes were centrifuged at 14,000 rpm for 5 minutes, and supernatant containing released [$^3$H]-oleic acid bound to BSA was collected and counted in a Tri-Carb liquid scintillation counter (PerkinElmer). The inhibitory effect of LA on cPLA2 was compared with that of AACOCF3 (Cayman Chemicals, #62120), PACOCF3 (Cayman Chemicals, #62650) and Pyrrolidine-2 (Calbiochem #525143) in the same assay conditions.

**Fig. S1**

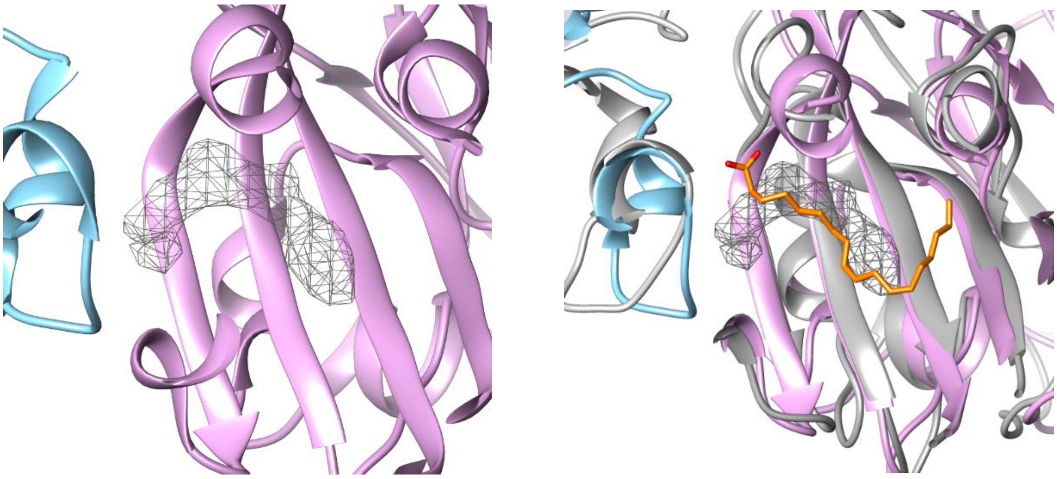

**Unassigned density in the cryo-EM structure of HCoV-OC43**. On the left, a zoomed view of the interface of two RBDs of HCoV-OC43 S is shown (PDB ID 6OHW). HCoV-OC43 RBDs are colored in blue and magenta. Unassigned density is shown as a mesh. A superimposition with the structure of locked SARS-CoV-2 S (PDB ID 6ZB5) in the same view is shown on the right. SARS-CoV-2 RBDs are colored in grey. LA is colored in orange, with oxygens of the polar head group colored in red.

**Fig. S2**

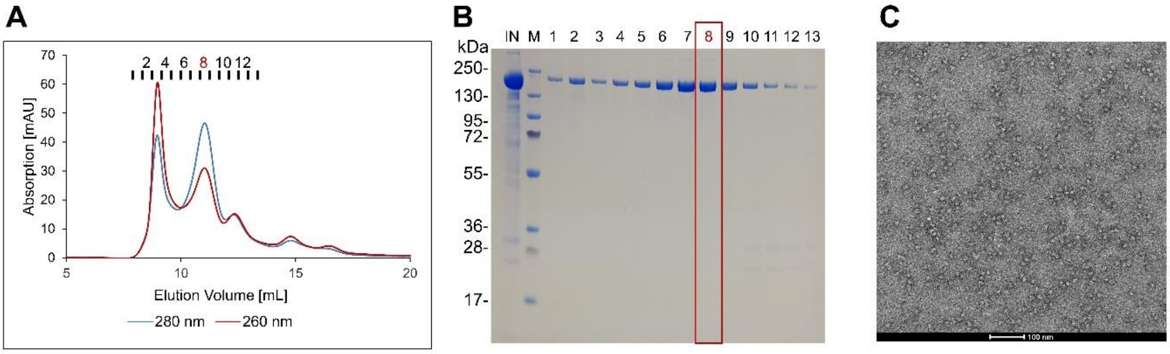


**Purification and quality control of SARS-CoV spike protein**. (**A**) Size-exclusion chromatogram of affinity-purified SARS-CoV spike protein using a Superdex 200 column. Absorption was detected at 280 nm (blue line) and 260 nm (red line). Peak fractions are indicated. The first peak at 8.6 mL elution volume corresponds to the void volume of the column comprising nucleic acids and spike protein as confirmed by the SDS-PAGE analysis in panel B (fractions 1 and 2). The spike trimer elutes at 11 mL. (**B**) SDS PAGE analysis of the SEC fractions from panel A. Lane 1: input fraction, lane 2: molecular weight marker, lane 3-15: fractions 1 to 13 from SEC. (**C**) Negative-stain EM micrograph of SEC peak fraction 8 (scale bar: 100 nm). In panels A and B peak fraction 8 is highlighted; this fraction was used for negative-stain EM and cryo-EM sample preparations.

**Fig. S3**

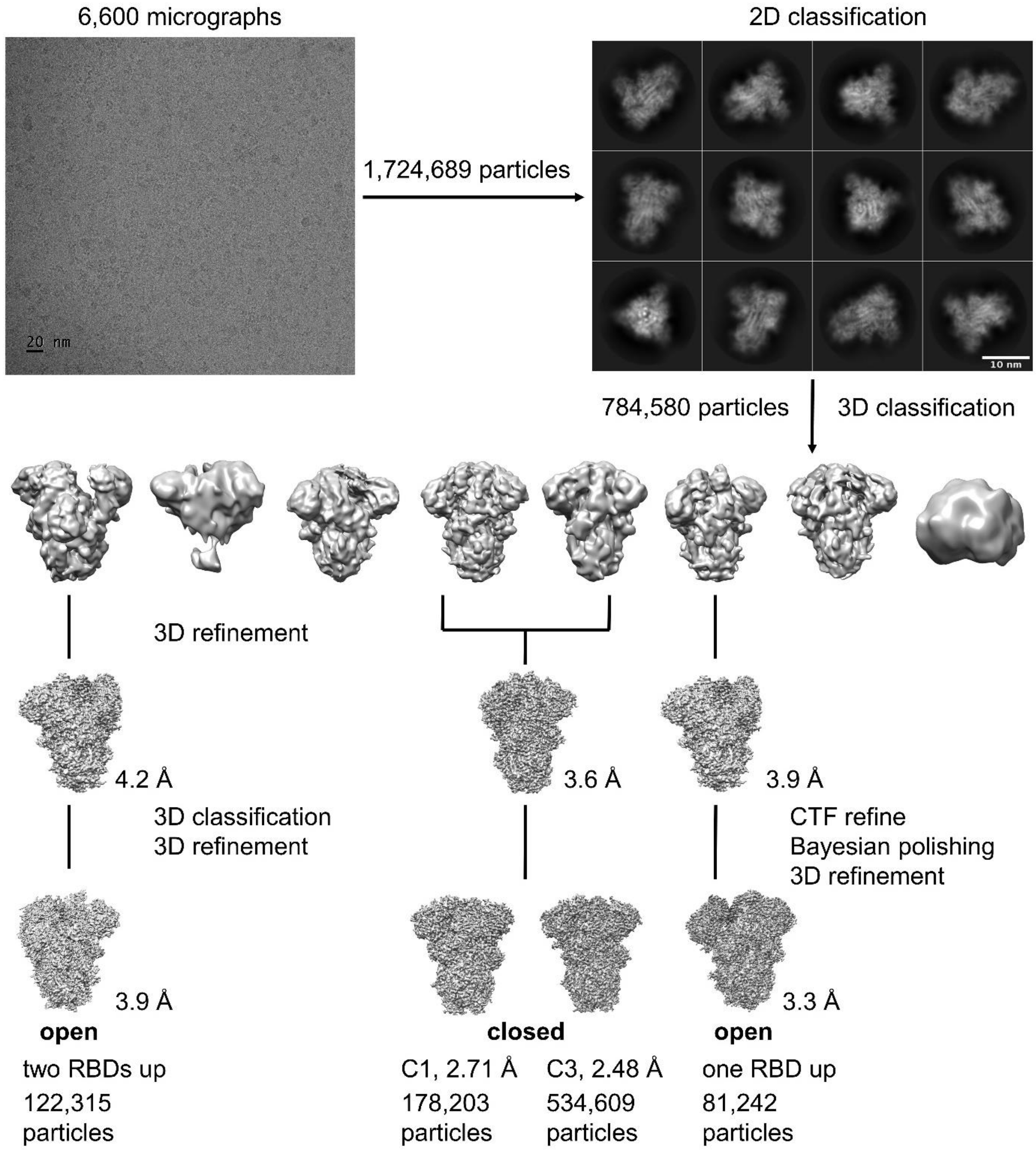


**Cryo-EM image processing workflow**. A motion-corrected cryo-EM micrograph (scale bar 20 nm), reference-free 2D class averages (scale bar 10 nm), 3D classification and refinement resulting in cryo-EM maps corresponding to the open conformations and the closed conformation (not symmetrized (C1) and C3-symmetrized) are shown.

**Fig. S4**

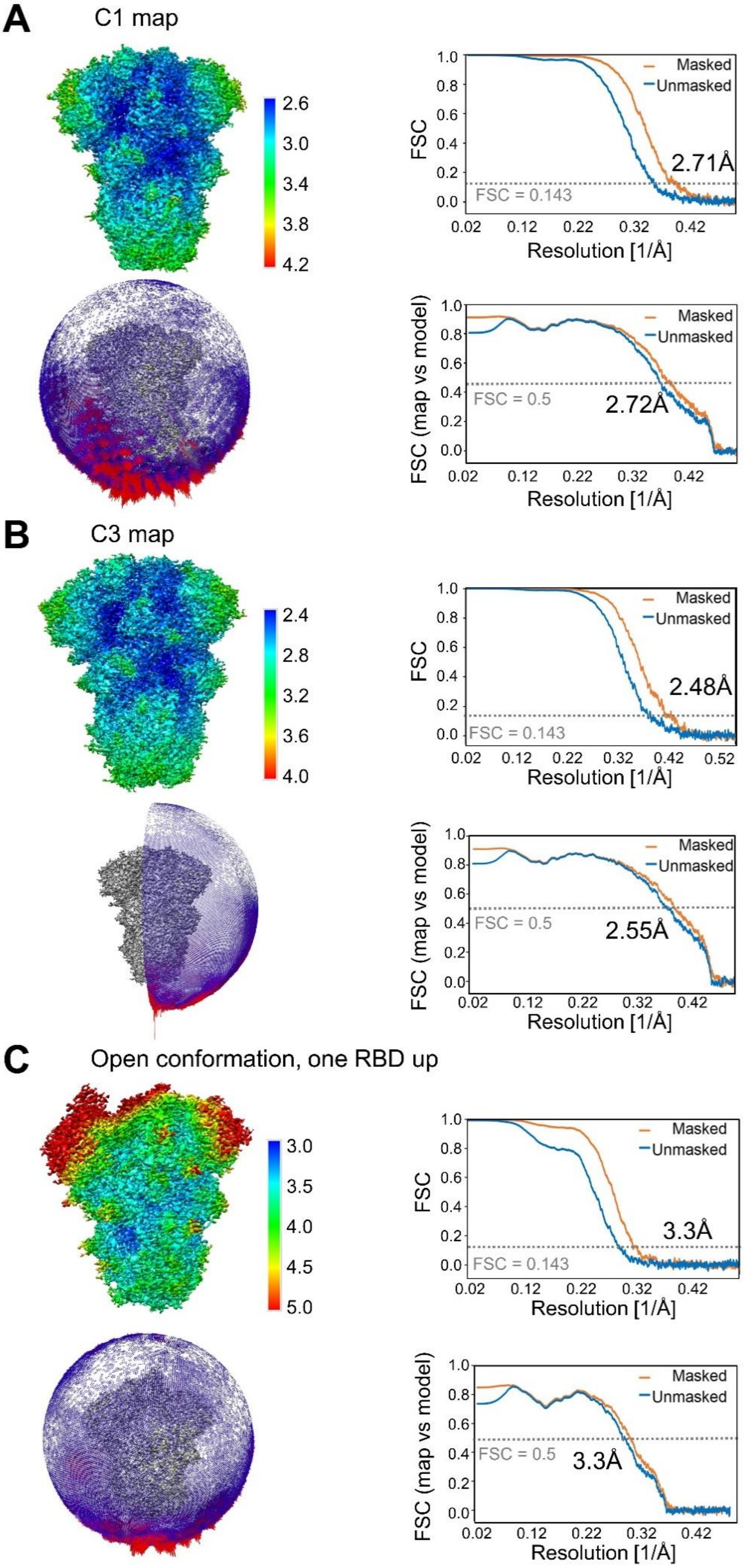
A
C1 map
2.6
3.0
3.4
3.8
4.2
FSC
Masked
Unmasked
2.71Å
FSC = 0.143
Resolution [1/Å]
FSC (map vs model)
FSC = 0.5
2.72Å
B
C3 map
2.4
2.8
3.2
3.6
4.0
2.48Å
2.55Å
C
Open conformation, one RBD up
3.0
3.5
4.0
4.5
5.0
3.3Å
3.3Å

**Cryo-EM structure validation.** Above left: Cryo-EM reconstruction colored according to the local resolution from a side view. Above right: Fourier Shell Correlation (FSC) curve after gold standard refinement. Below left: Orientation distribution of views that contributed to this map. Longer red rods represent orientations that comprise more particles. Below right: Cross-validation FSC curves for the refined model versus the final masked and unmasked maps. Corresponding panels are shown for (**A**) the closed unsymmetrized C1 map (**B**) the closed C3-symmetrized map and (**C**) the open, one RBD up conformation map.

**Fig. S5**

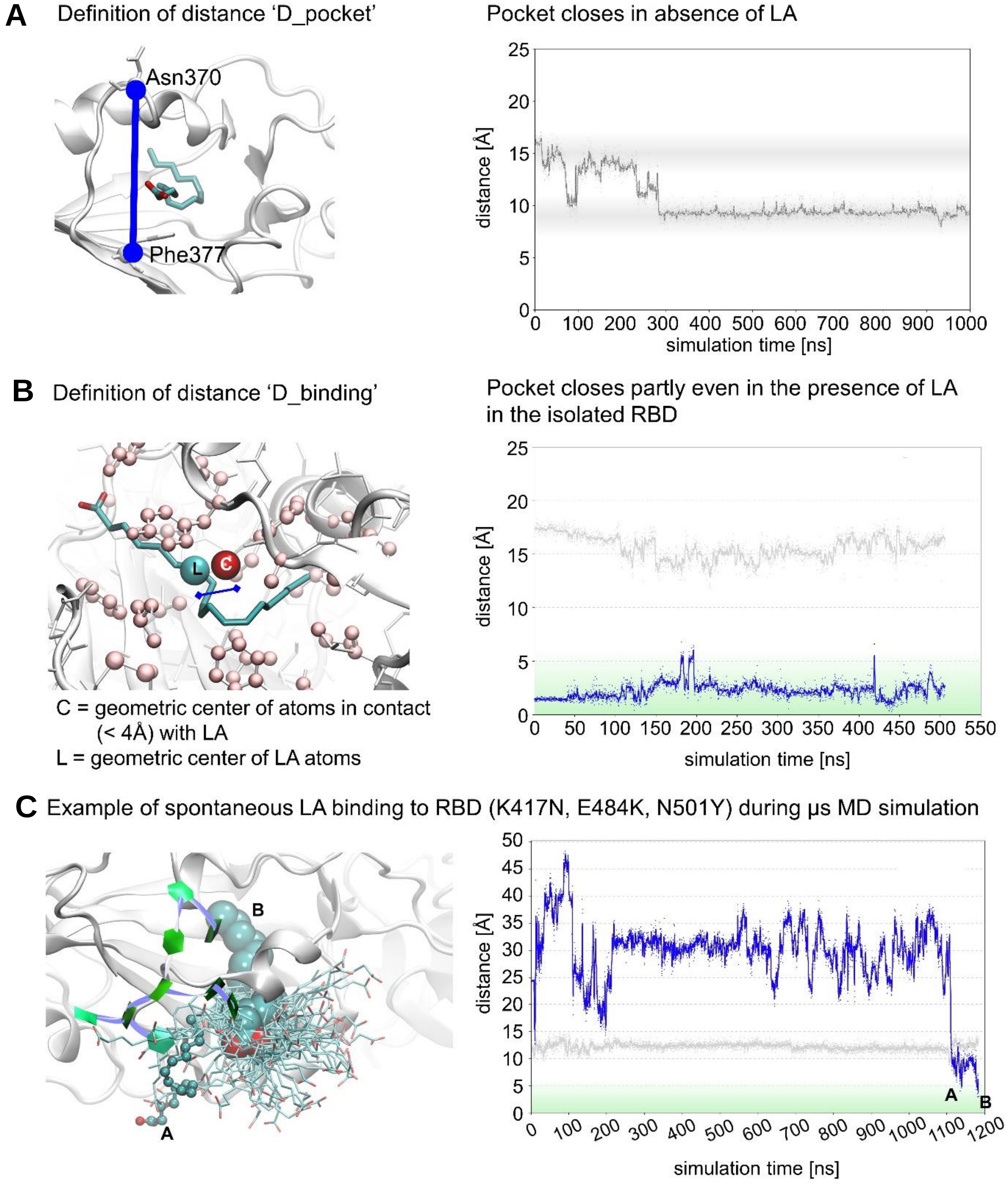


**MD analysis of pocket dynamics and LA-binding to SARS-CoV-2 spike RBD.** (**A**) D_pocket (grey curve, right) is the distance between the Cα atoms of residues Asn370 and Phe377) shown as blue spheres in the RBD structure on the left (PDB ID 6ZGE (*58*)). D_pocket thus monitors the dynamic opening and closing of the pocket, as visualized on the right side. Open and closed states are indicated by gray bars at ~15 Å and ~9 Å, respectively. After eliminating LA, the pocket closes, indicated by a decrease of D_pocket as shown on the right.

(**B**) LA dynamics measured by D_binding. C is the geometric center of the pocket. Left panel: D_binding is the distance between the center of the LA molecule (L) and C. L and C are shown as cyan and red spheres in the RBD structure (PDB ID 6ZGE). Right panel: Plotted are both D_binding (blue curve) and D_pocket (grey curve), monitoring the binding and dynamics of LA in the pocket (or in the case of hCoV-HKU1 to a hydrophobic site on the surface of the RBD) and pocket opening, respectively. The green bar indicates distances compatible with LA-binding in the pocket. In the isolated LA-bound RDB, the pocket entrance, as measured by D_pocket (grey curve), appears to be slightly smaller than in the locked spike trimer. (**C**) An additional successful binding trajectory is shown for the B.1.351/ beta variant (see also Fig 3B). Here, the pocket was closed in the starting structure. Left panel: The binding event (time between A and B) is represented as a sequence of overlayed MD snapshots to show that LA entry into the pocket is 'highly dynamic'. LA is shown in teal stick representations, with time point A as small spheres, time point B as large spheres, and all other snapshots as stick only and with oxygens of the polar head group colored in red. Right panel: D_pocket (grey curve) and D_binding (blue curve) distances are shown.

**Fig. S6**

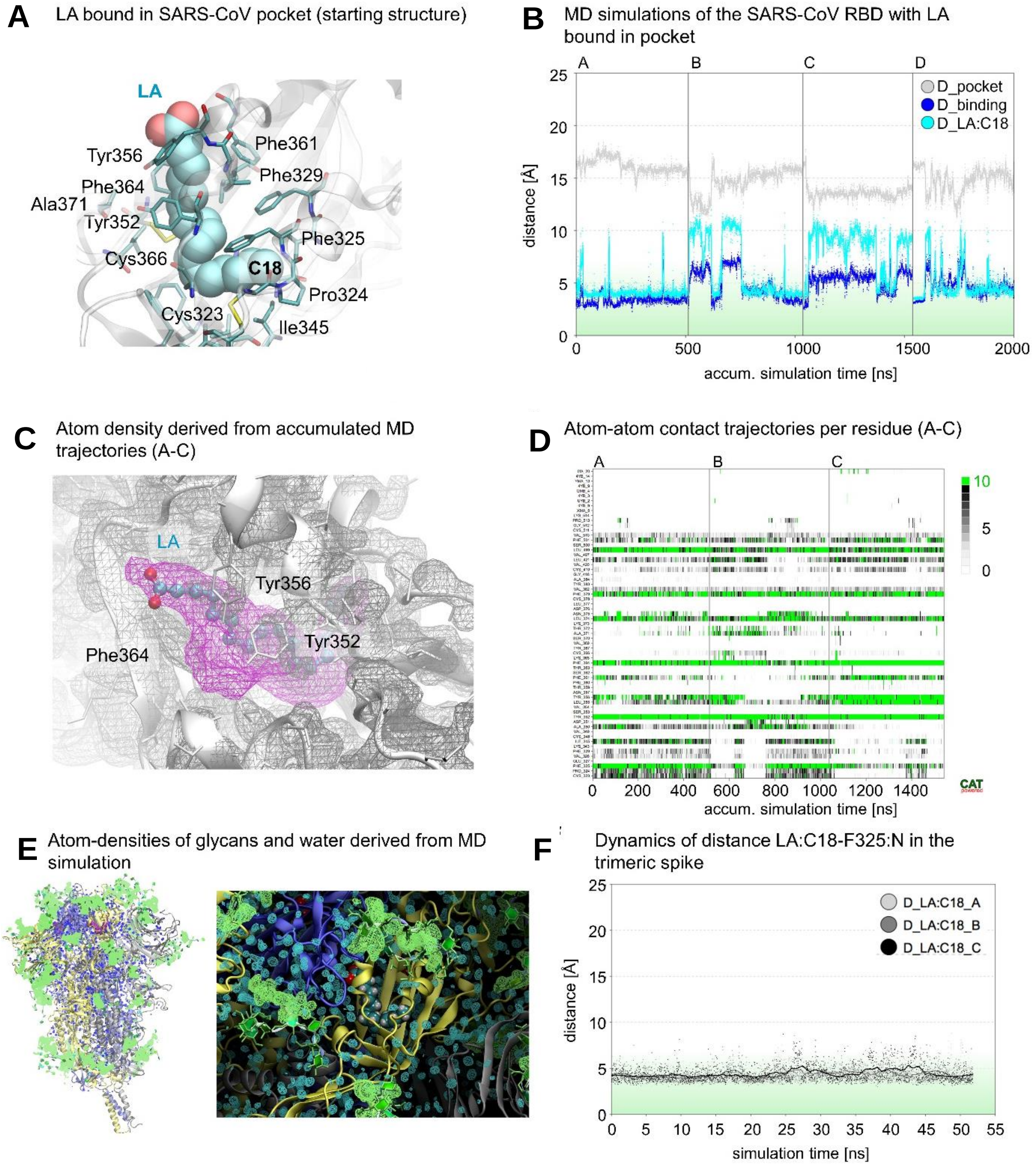


**MD analysis of dynamic LA-binding to SARS-CoV spike RBD.** (**A**) Starting SARS-CoV spike RBD structure with LA bound in the pocket (this study). Hydrophobic residues forming the pocket and the C18 atom of LA are labeled. (**B**) 2 µs MD analysis with LA 'stably' bound in the pocket of isolated RBD. However, the C18 atom of LA (cyan curve) can move away from the sub-pocket formed by Phe325, Pro324, Ile345 and Cys323 (see structure in panel A). Trajectories for D-pocket (grey curve), D_binding (blue curve) and D_LA:C18 (cyan curve, being a measure of the distance between the C18 atom of LA and the geometric center of the pocket) are plotted. (**C**) The dynamic LA-binding in the pocket is shown as density (pink mesh) derived from the accumulated MD trajectories A-C shown in panel B. (**D**) Number of atom-atom contacts (<4 Å) of residues forming the LA-binding pocket (Y-axis) with LA, derived from the MD trajectories A-C in panel B. In green (corresponding to 10 atom-atom contacts or more) the dynamics of strong interactions are highlighted, *e.g.,* with Phe325, Tyr352, Tyr356, Phe364, Leu374, Phe379 and Leu499. (**E**) Analysis of LA-binding to SARS-CoV spike. Left panel: A side view of a single MD snapshot of the SARS-CoV spike (protein chains are depicted in grey, yellow and blue cartoon representation). Right panel: A zoomed view of the LA-binding pocket. LA represented as cyan spheres with hydrogens and carbons in white and red respectively and glycans as sticks in light green. In the closed/locked SARS-CoV spike trimer, LA appears rigidly positioned in a single binding mode. (**F**) Analysis of the dynamics of LA in the pocket (D_LA:C18). The distance LA:C18-F325:N is shown in 50 ns MD simulations of the trimeric spike for the three LA-pockets (in RBD chains A, B and C), confirming that LA is firmly bound in all three pockets. D_pocket and D_binding are defined in fig. S5A,B.

**Fig. S7**

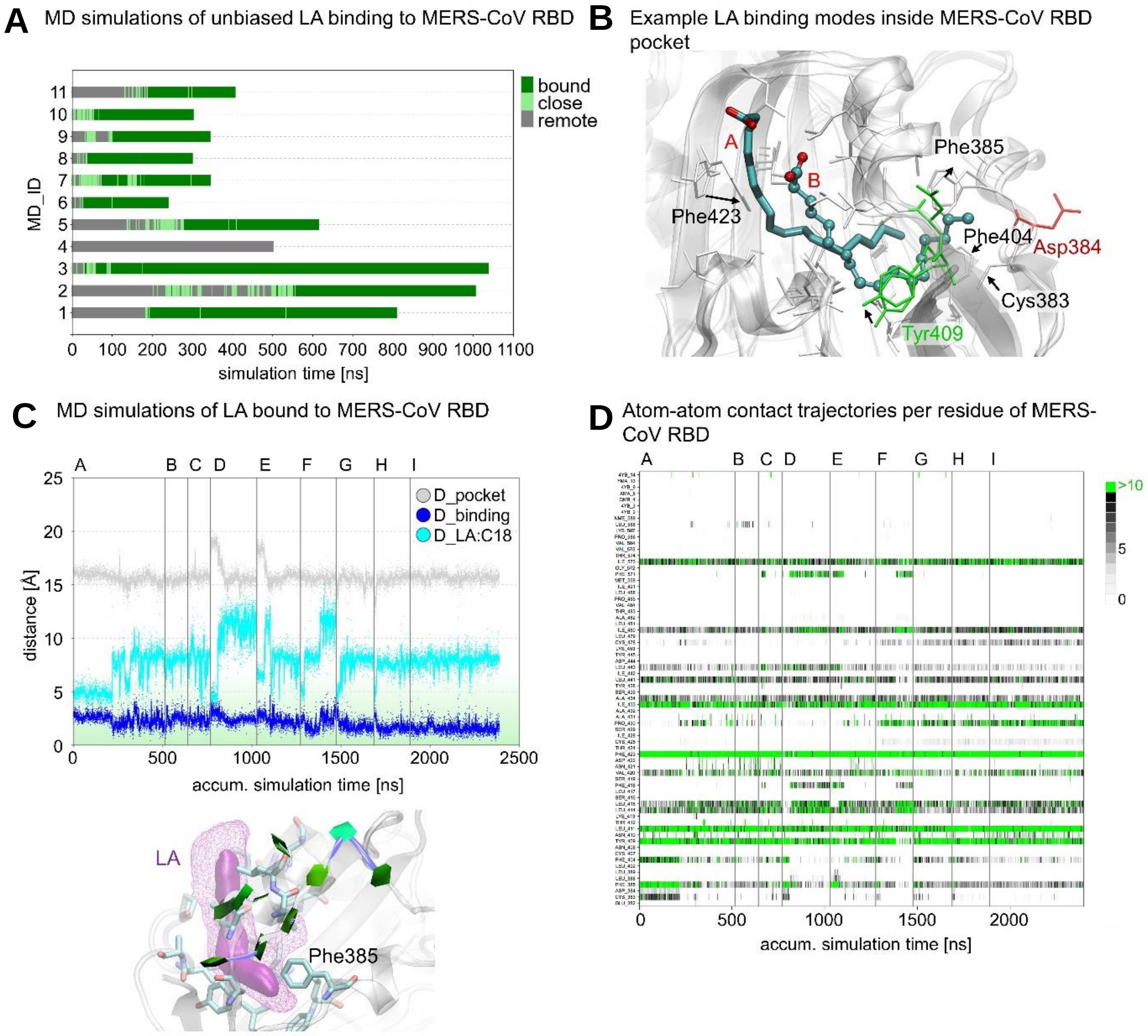


**LA-binding to the pocket in the RBD of MERS-CoV spike.** (**A**) 11 MD trajectories are shown for LA-binding to the pocket in MERS-CoV RBD. Bound (LA in the pocket), close (LA at pocket entrance) and remote states are shown in green, light green and grey, respectively. In 10 simulations, spontaneous LA-binding (green) was observed in less than 500 ns. (**B**) Following the MD simulations, LA is binding in the pocket mainly in mode A (with a distance D_LA:C18 of about 8 Å, LA is represented as cyan sticks). However, in analogy to SARS-CoV spike MD simulations (fig. S6), a binding mode B (with C18 of LA bound to a pocket formed by Phe385, Asp384, Cys383 and Phe404, LA is represented as cyan spheres) may be also plausible. (**C**) Above: Multiple MD trajectories of LA bound to the MERS-CoV RBD. The different distances D_pocket (grey curve), D_binding (blue curve) and D_LA:C18 (cyan curve) are shown as accumulated trajectory plot. Below: the LA atom density calculated from the accumulated MDs is shown as iso-contour plot in pink (solid: high atom density; mesh:

lower atom density) within the MERS-CoV spike RBD. LA-binding inside the pocket is stable. However, the short distance (<5Å) between LA:C18 and F385:N is not maintained after the restraints are released. (**D**) Number of atom-atom contacts (<4 Å) of LA with residues forming the LA-binding pocket (Y-axis) in the MERS-CoV spike RBD, derived from MD simulations (panel C). Strong interactions (10 or more contacts) are depicted in green.

**Fig. S8**

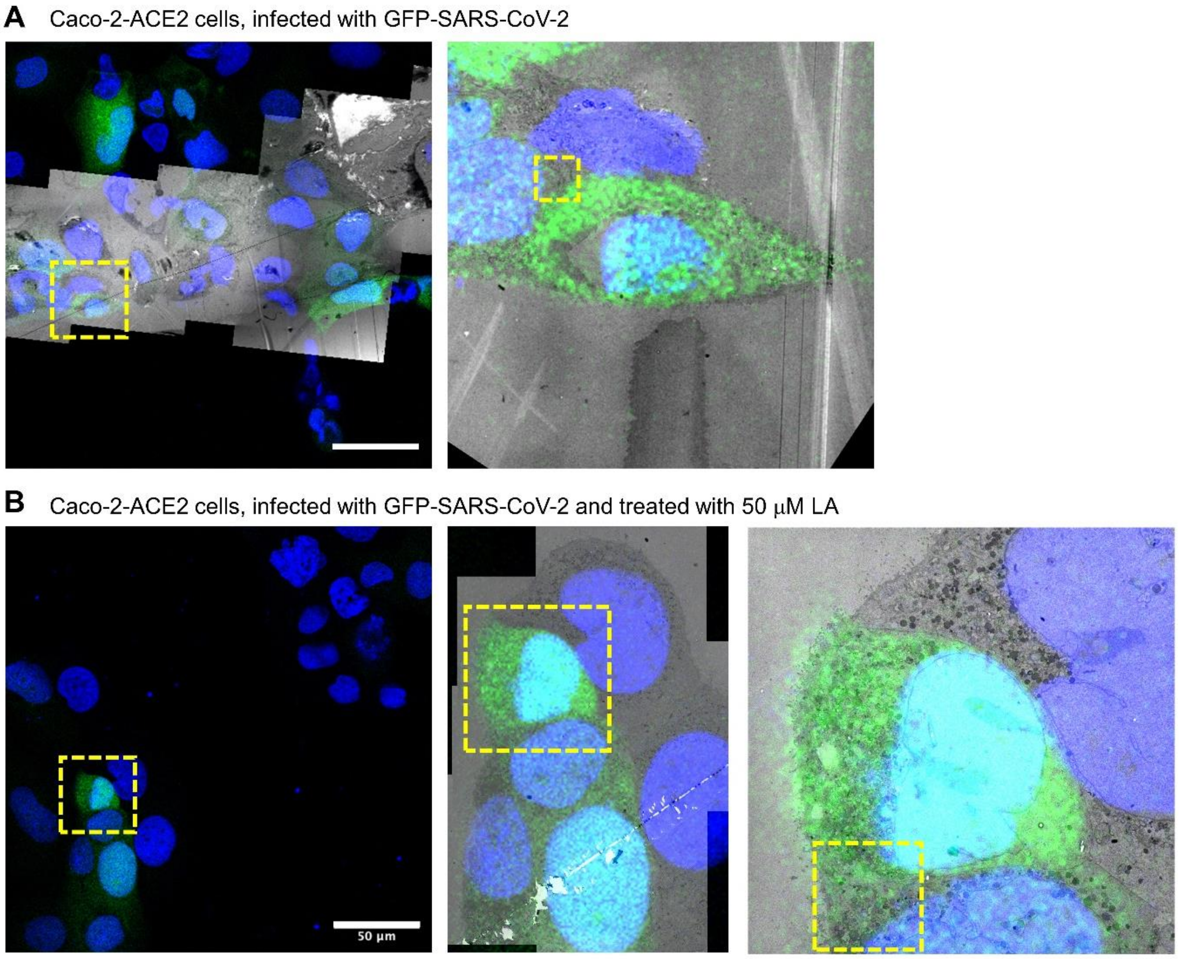


**Correlative Light-Electron Microscopy (CLEM) of GFP-expressing SARS-CoV-2 infected Caco-2-ACE2 cells.** (**A**) Overlay of fluorescence microscopy image and TEM mosaic. Left: overview image with a scale bar of 50 μm. The yellow box highlights the region of interest used for further analysis. Right: Close-up view of the cells chosen for further analysis. The yellow box highlights the region shown in Fig. 4A which was used for electron tomography. Blue indicates DAPI staining of nuclei and green the GFP-SARS-CoV-2 virus. (**B**) Left: overview image from fluorescence microscopy for Caco-2-ACE2 cells infected with GFP-SARS-CoV-2 virus (green) and treated with 50 μM LA 1-hour after infection. A green infected cell was chosen for further analysis. Middle and right image: Close-up views of the SARS-CoV-2 infected cell. The yellow box (left image) highlights the region shown in Fig. 4B which was used for electron tomography.

**Fig. S9**

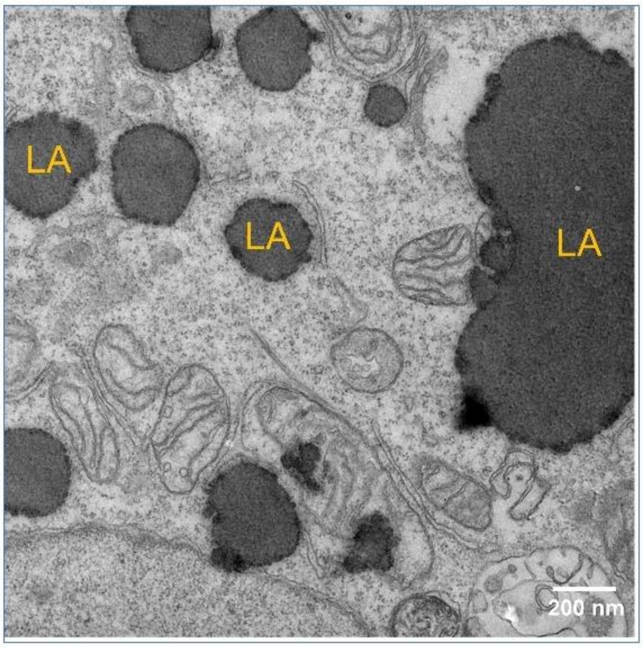


**Uninfected Caco-2-ACE2 cell treated with LA.** LA is readily taken up by the cells and appears as high contrast regions (marked with LA). The scale bar (200 nm) is colored in white.

**Fig. S10**

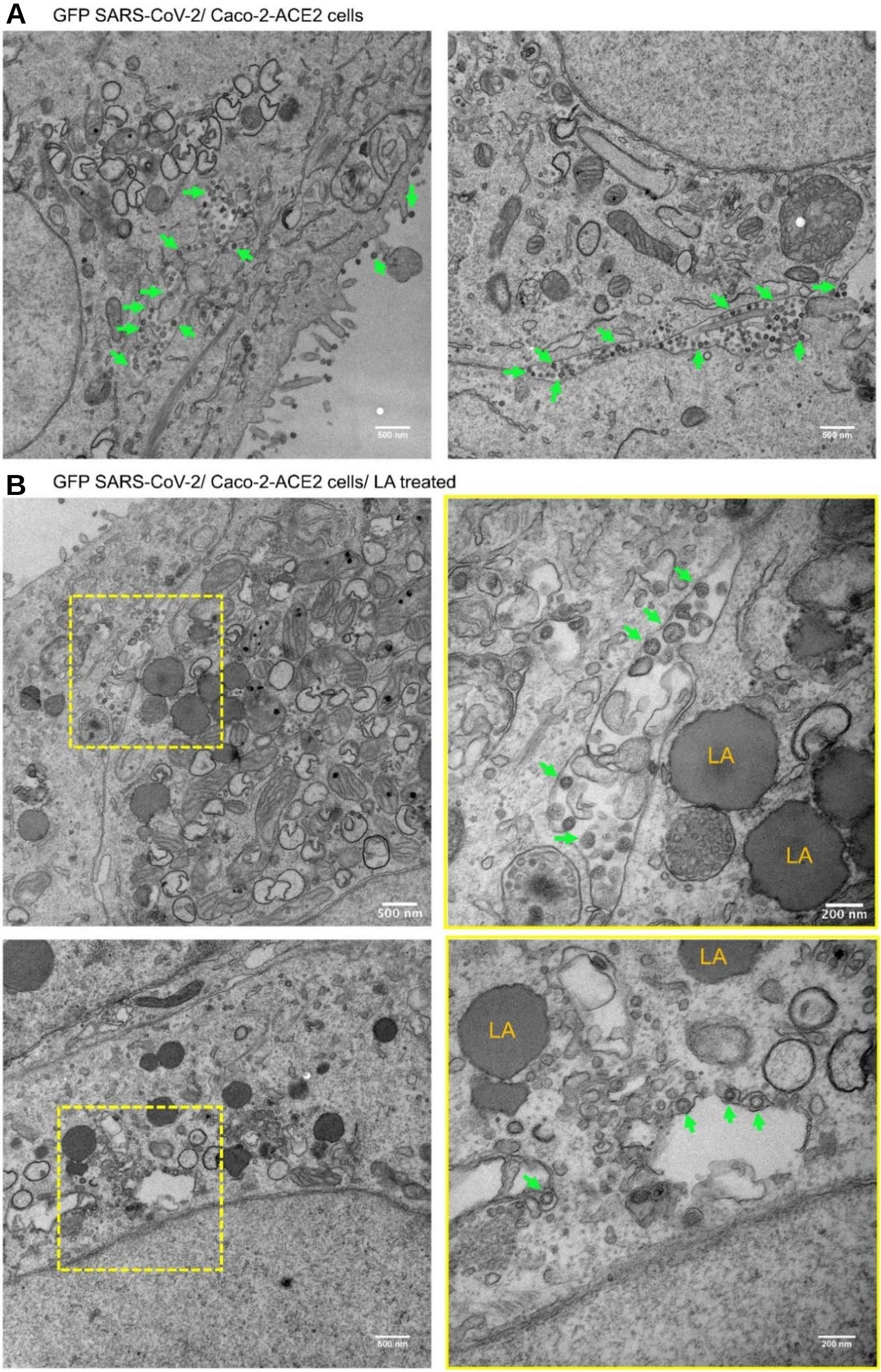
A
GFP SARS-CoV-2/ Caco-2-ACE2 cells
500 nm
500 nm
B
GFP SARS-CoV-2/ Caco-2-ACE2 cells/ LA treated
500 nm
LA
LA
200 nm
500 nm
LA
LA
200 nm

**EM analysis of GFP-expressing SARS-CoV-2 infected Caco-2-ACE2 cells.** Only green (GFP-expressing) cells were analyzed. Scale bar: 500 nm. (**A**) Image of two different cells in the absence of LA treatment. Green arrows point to virions. (**B**) Left: Overview images of GFP-SARS-CoV-2 infected cells treated with 50 μM LA after infection. Scale bar: 500 nm. Yellow boxes highlight regions with virions. Right: Close-up views. Scale bar: 200 nm. Green arrows point to virions. LA highlights lipid droplets.

**Table S1**

**LA binding constants $K_D$, $k_{on}$ and $k_{off}$.**

| **β-CoV S, RBD** | **$K_D$, LA (nM)** | **$k_{on}$ ($M^{-1}s^{-1}$)** | **$k_{off}$ ($s^{-1}$)** |
|---|---|---|---|
| **SARS-CoV-2 *** | 41.1±16 | 1.0±0.4x$10^4$ | 3.9±1.3 x$10^{-4}$ |
| **SARS-CoV-2 alpha** | 50±15.2 | 9.2±6.5 x$10^4$ | 4.1±2.0 x$10^{-3}$ |
| **SARS-CoV-2 beta** | 86.3±10.5 | 9.3±3.7 x$10^4$ | 8.3±3.8 x$10^{-3}$ |
| **SARS-CoV-2 gamma** | 87.0±8.5 | 5.4±2.2 x$10^3$ | 4.7±1.7 x$10^{-4}$ |
| **SARS-CoV-2 delta** | 35.2±25.0 | 3.2±1.5 x$10^4$ | 0.9±0.4 xx$10^{-3}$ |
| **SARS-CoV-2 omicron** | 39.6±11.9 | 1.1±0.4 x$10^4$ | 0.5± 0.3x$10^{-3}$ |
| **SARS-CoV** | 72 ± 36 | 2.4±0.6 x$10^4$ | 1.7±1.0 x$10^{-3}$ |
| **MERS-CoV** | 96 ± 35 | 3.0±1.6 x$10^4$ | 2.5±0.6 x$10^{-3}$ |
| **HCoV-HKU1** | - | - | - |
| **HCoV-HKU1 E375A** | 178 ± 37 | 1.8±0.6 x$10^4$ | 3.0±0.5 x$10^{-3}$ |

**Value reported in Ref (11).*

**Table S2**

**Cryo-EM data collection and refinement statistics for SARS-CoV.**

| | **Closed conformation, C3 symmetrized** | **Closed conformation, C1** | **Open conformation, C1** |
|---|---|---|---|
| Voltage (kV) | 200 | 200 | 200 |
| Magnification (nominal) | 130,000 | 130,000 | 130,000 |
| Pixel size (Å/pix) | 1.05 (0.525) | 1.05 (0.525) | 1.05 (0.525) |
| Flux ($e^-$ /pix/sec) | 5.77 | 5.77 | 5.77 |
| Frames per exposure | 60 | 60 | 60 |
| Exposure ($e^-$ /Å$^2$) | 1.04 | 1.04 | 1.04 |
| Defocus range (μm) | -0.8 to -2.0 | -0.8 to -2.0 | -0.8 to -2.0 |
| Micrographs collected | 6603 | 6603 | 6603 |
| Particles, final | 534,609 | 178,203 | 81,242 |
| Map sharpening B-factor (Å$^2$) | -73.71 | -68.0 | -100.8 |
| Masked resolution at 0.143 FSC (Å) | 2.48 Å | 2.71 Å | 3.3 Å |

**Refinement**

| | **Closed conformation, C3 symmetrized** | **Closed conformation, C1** | **Open conformation, C1** |
|---|---|---|---|
| Composition | | | |
| Amino acids | 3060 | 3060 | 2785 |
| Glycans | 36 | 33 | 25 |
| Ligands | 3 | 3 | - |
| RMSD bonds (Å) | 0.004 | 0.004 | 0.006 |
| RMSD angles (º) | 0.617 | 0.586 | 0.671 |
| Mean B-factors (Å$^2$) | | | |
| Amino acids | 30.44 | 22.76 | 25.56 |
| Ligands | 28.52 | 29.97 | 40.26 |
| Ramachandran | | | |
| Favored (%) | 96.32 | 96.29 | 89.15 |
| Allowed (%) | 3.48 | 3.68 | 10.74 |
| Outliers (%) | 0.2 | 0.03 | 0.11 |
| Rotamer outliers (%) | 2.19 | 2.35 | 4.49 |
| Clash score | 2.28 | 2.85 | 5.65 |
| C-beta outliers (%) | 0.00 | 0 | 0.00 |
| CaBLAM outliers (%) | 3.66 | 3.53 | 6.37 |
| CC (mask) | 0.79 | 0.81 | 0.76 |
| MolProbity score | 1.51 | 1.60 | 2.37 |
| EMRinger score | 3.85 | 3.82 | 2.5 |
| Model resolution (Å) 0.5 FSC threshold | 2.6 | 2.9 | 3.3 |

### Table S3
### N-linked glycosylation sites in the SARS-CoV spike

| SARS-CoV spike | |
|---|---|
| **WT* predicted** | **Recombinant, expressed in Hi5** |
| **UNIPROT (P59594)** | **this study **** |
| $N_{29}$YT | |
| $N_{65}$VT | $N_{65}$VT |
| $N_{73}$HT | |
| $N_{109}$KS | $N_{109}$KS |
| $N_{118}$NS | |
| $N_{119}$ST | $N_{119}$ST |
| $N_{158}$CT | $N_{158}$CT |
| $N_{227}$IT | $N_{227}$IT |
| $N_{269}$GT | $N_{269}$GT |
| $N_{318}$IT | $N_{318}$IT |
| $N_{330}$AT | $N_{330}$AT |
| $N_{357}$ST | $N_{357}$ST |
| $N_{589}$AS | |
| $N_{602}$CT | $N_{602}$CT |
| $N_{691}$NT | $N_{691}$NT |
| $N_{699}$FS | $N_{699}$FS |
| $N_{783}$FS | $N_{783}$FS |
| $N_{1056}$FT | $N_{1056}$FT |
| $N_{1080}$GT | $N_{1080}$GT |
| $N_{1116}$NT | |
| $N_{1140}$HT | |
| $N_{1155}$AS | |
| $N_{1176}$ES | |
| ***** *NC_004718.3 (Isolate BJ01) (SARS-CoV)*<br>***Sites lacking glycosylation in cryo-EM maps are omitted (boxes colored in grey)* | |

**Movies Captions**

**Movie S1:** Simulation of LA-binding to the SARS-CoV-2 RBD (PDB ID 6zb5 (*11*)) viewed from the pocket entrance. LA is shown as spheres colored teal for carbon and red for the oxygen in the carboxyl headgroup.

**Movie S2:** Simulation of LA-binding to the pocket in SARS-CoV RBD (this study), viewed from the pocket entrance and then from inside the pocket. LA is shown as spheres colored teal for carbon and red for the oxygen in the carboxyl headgroup.

**Movie S3:** Simulation of LA-binding to the pocket in MERS-CoV RBD (PDB ID 6q05 (*48*), with closed pocket), viewed from the pocket entrance. LA is shown as spheres colored teal for carbon and red for the oxygen in the carboxyl headgroup.

**Movie S4:** Simulation of LA-binding to the pocket in the SARS-CoV-2 omicron RBD (from PDB ID 7oaq (*59*), with closed pocket), viewed from the pocket entrance. LA is shown as spheres colored teal for carbon and red for the oxygen in the carboxyl headgroup.

**Movie S5:** Digital sections through an electron tomogram of a SARS-CoV-2 infected Caco-2-ACE2 cell at 36 hours after infection, as shown in Fig. 4A. To facilitate image alignment during image reconstruction, a suspension of 15-nm gold particles was layered on each side of the sections as fiducial markers.

**Movie S6:** Digital sections through an electron tomogram of a SARS-CoV-2 infected Caco-2-ACE2 cell at 36 hours after infection, treated with 50 μM LA 1-hour after infection, as shown in Fig. 4B. To facilitate image alignment during image reconstruction, a suspension of 15-nm gold particles was layered on each side of the sections as fiducial markers.
